# “Integration of multimodal data in the developing tooth reveals candidate dental disease genes”

**DOI:** 10.1101/2022.03.15.483501

**Authors:** Emma Wentworth Winchester, Alexis Hardy, Justin Cotney

## Abstract

Dental malformations range from rare syndromes to common nonsyndromic phenotypes. These malformations can predispose individuals to dental disease, which can in turn affect systemic health. While many dental phenotypes are heritable, most cases have not been linked to deleterious mutations in single genes. We demonstrate that human and conserved mouse craniofacial enhancers show enrichment of dental phenotype-associated variants. Given these findings in bulk craniofacial tissues, we looked to determine the role of tooth enhancers in this phenomenon. We used ChIP-seq and machine learning to identify enhancers of E13.5 mouse incisors. Multi-tissue comparisons of human and mouse enhancers revealed that putative tooth enhancers had the strongest enrichment of dental phenotype-associated variants, suggesting a role for dysregulation of tooth development in dental phenotypes. To uncover novel dental phenotype-driving genes in the developing tooth we performed coexpression analysis and annotated the contributing cell types of gene modules using scRNAseq. Through integration of chromatin state, bulk gene coexpression, and cell type resolved gene expression we prioritized a list of candidate novel dental disease genes for future investigations in mouse models and human studies.

## Background

Dental malformations range from rare syndromes such as amelogenesis or dentinogenesis imperfecta [1–5], to relatively common nonsyndromic phenotypes, such as abnormal tooth number [6–10] or enamel hypoplasia [11,12]. Aberrations from normal dental development may predispose individuals to dental disease [13,14] and require costly and complex interventions. As dental health is known to impact systemic health [15,16], dental malformations pose a substantial public health burden and determining their causes is a priority. While environmental factors play a major role in the incidence of many of these common phenotypes, the heritability of normal and abnormal dental phenotypes has been reported to range from 40 to 90% suggesting strong genetic components [17–23].

In most mammals the primary teeth begin to develop during early embryogenesis and are mostly developed before erupting into the oral cavity shortly after birth. As such, the underlying genetic causes of many dental malformations and diseases likely exert their effects during development. Therefore, genetic programs that coordinate dental development are of particular interest in identifying drivers of dental malformations and disease. While rare syndromes affecting teeth have been linked to mutations in specific genes [24–27], most cases of common dental malformations, diseases, and phenotypes such as caries [20–23,28,29], delayed tooth eruption [30], and abnormal tooth number [31–33] have not been linked to deleterious mutations in single genes or large DNA copy number changes. The combination of relatively high heritability of many of these dental phenotypes despite only a small fraction of cases being explained by mutations in protein-coding genes and their tooth-isolated nature suggests that these phenotypes may be associated with defective gene regulation caused by enhancers.

Enhancers are gene regulatory sequences that can operate over long genomic distances, some over 1 megabase from their target gene, to regulate gene expression in spatiotemporal-specific contexts. [34–36] Disruption of enhancers results in changes in gene expression when and where that enhancer is normally active, resulting in typically tissue-isolated abnormalities. [37,38] This concept has been shown to contribute to craniofacial phenotypes such as nonsyndromic orofacial clefting and normal craniofacial morphology. [39–41] Common human single nucleotide polymorphisms (variants) associated with these craniofacial phenotypes are enriched in enhancer sequences active in embryonic human craniofacial tissues. However, findings in the bulk human face have not been extended to investigate the contribution of these enhancers in dental morphogenesis or disease. As a consequence of this, the target genes of any dental phenotype-associated enhancers that ultimately drive the phenotype have not been identified in a systematic manner.

Ideally investigations of these drivers would use primary developing human teeth. However, due to the gestational time period in human development when these cell types arise [42–46] and lack of clear *in vitro* models, tooth development is primarily studied using primary mouse tissue. The early stages of development of the mouse dentition are largely analogous to human dental development [42–44], and many rare human dental syndromes are modelable in mice. [42,44,47–49] However, investigations into the noncoding genome of the developing tooth in mammals have been limited to a small number of regions [50,51]. As such, the conservation of the noncoding drivers of dental development between human and mouse genome-wide has not been definitively proven.

In this manuscript we demonstrate the contribution of active human and mouse craniofacial enhancers to dental disease risk and dental developmental traits. We used ChIP-seq and a well-established chromatin annotation pipeline [40,52,53] to annotate active enhancers of the developing tooth and demonstrate that these regions have the highest burden of dental phenotype-associated variants compared to all other tissues. To uncover potential disease-causing target genes associated with these enhancers, we integrated bulk and single cell transcriptomics with our ChIP-seq enhancer calls. Specifically, we leveraged weighted gene co-expression network analysis (WGCNA) of publicly available bulk transcriptomic data whereby we identified modules of coexpressed genes specifically relevant to the developing tooth. We determined the cell type specificity of these modules using novel cell type specific transcriptomic signatures identified from re-analysis of publicly available scRNA-seq data of the E14 mouse molar. Transcriptomic data were integrated with E13.5 mouse incisor enhancer calls and human Genome-Wide Association Study data (GWAS), which were used to prioritize candidate disease genes. We present the results of this analysis as a prioritized list of candidate human disease genes with the intention of their use in future in-depth experiments. In addition to these genes, the tissue-specific enhancers we describe are of particular interest as they provide precise molecular tools for tissue specific knockouts or reporter gene expression as we have described for specific regions of the mouse brain [54].

## Results

### Section 1. Craniofacial enhancers contribute to dental development and disease risk

It has been well described that the regulatory regions active in early craniofacial development play an important role in orofacial clefting and normal human facial variation [39–41,55]. Given the demonstrated genetic component of nonsyndromic dental phenotypes such delayed tooth eruption and hypodontia, we hypothesized that craniofacial regulatory regions may also play a role in these relatively common phenotypes. To test this, we compiled a list of ‘odontogenesis’ and ‘caries’-associated common variants from the GWAS catalog (GO_0042476 and EFO_0003819, respectively; Supplemental Table 1)[56–62]. We interrogated the enrichment of these dental phenotype-associated variants (DVs) within strong enhancers of the developing human face [40] compared to strong enhancers of hundreds of other tissues [52](Methods). We observed an enrichment of variants associated with dental phenotypes in active enhancers of craniofacial tissues compared to hundreds of other tissue types from the Roadmap Epigenome Project, for variants associated with both odontogenesis (Figure 1A) and caries (Figure 1B). Notably, enrichment is not observed in craniofacial enhancers for variants associated with diseases not known to have a craniofacial component, such as Crohn’s disease (EFO_0000384)(Supplemental Figure 1). These results suggest a role for the regulatory regions active in the developing craniofacial apparatus in dental developmental traits and disease.

**Figure 1.**
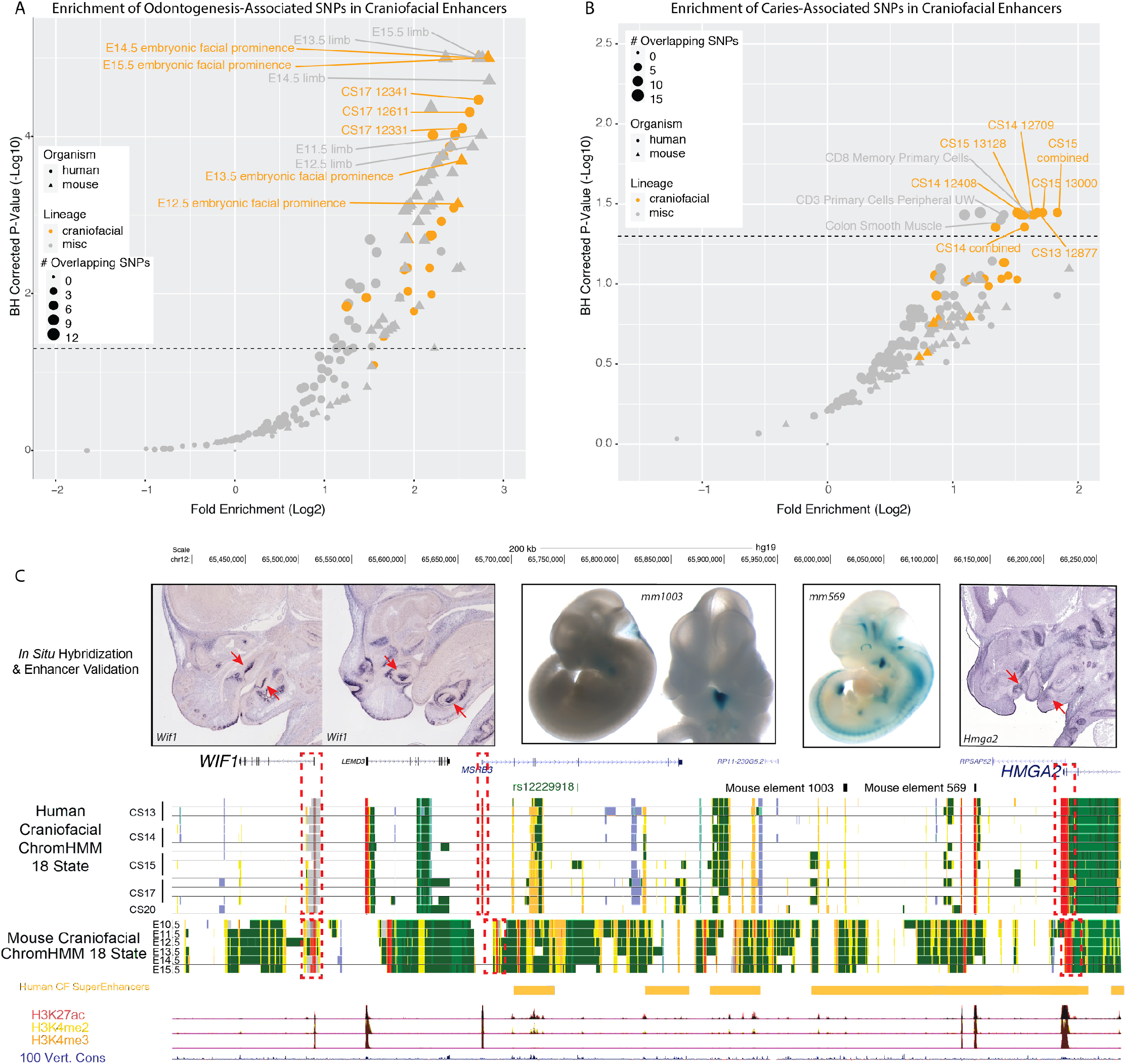
Conserved Craniofacial Enhancers Contribute to Dental Development and Phenotype Risk A. Scatterplot of GREGOR (genomic regulatory elements and GWAS overlap algoRithm) analysis of enrichment of “Odontogenesis” (GO_0042476)-associated variants from GWAS Catalog in active enhancers annotated by 18 state chromatin segmentations genome-wide for samples from [40](orange circles), Roadmap Epigenome (grey circles), and 18-state segmentations from Mouse ENCODE craniofacial (orange triangles) and other mouse tissue samples (grey triangles) [40,52,63]. B. GREGOR [64] analysis of “Caries” (EFO_0003819)-associated variants from GWAS Catalog as in A. C. Visualization of the locus surrounding rs12229918 (chr5:134509987, hg19) and nearby fine-mapped variants associated with decreased tooth number and delayed tooth eruption by [57]. Mouse craniofacial 18 state segmentations were generated as described in Methods from data from ENCODE [63]. Human craniofacial 18 state segmentation and composite H3K27ac, H3K4me2, and H3K4me3 p-value signal tracks were obtained from [40]. *In situ* hybridization images of *Wif1* and *HMGA2* were obtained from GenePaint [65]. Enhancer assay images were obtained from VISTA [66].

These findings raised the question whether loci of risk for dental phenotypes are similarly conserved in the widely used developmental model, the mouse. To determine this, we interrogated enrichment of the same variants in conserved strong enhancers from hundreds of publicly available embryonic and postnatal mouse samples, including the developing craniofacial structures (Methods)[63]. Similar to humans, we observed an enrichment of odontogenesis (Figure 1A) associated variants in conserved mouse craniofacial active enhancers. When we interrogated caries-associated variants (Figure 1B), no mouse tissue passed genome-wide significance cutoff. The combination of these findings in human and mouse support our hypothesis that some common dental phenotypes, specifically age of tooth eruption and number of teeth, may be directly impacted by distal gene regulatory sequences and that these regions may affect generally conserved processes in orofacial development.

While these results indicated a general role in craniofacial enhancers and development in dental phenotypes, it did not initially reveal any particular target genes important for these phenotypes. When we inspected the loci implicated by our analysis, we observed a variant associated with decreased tooth number (rs12229918) located in an intron of *MSRB3. MSRB3* does not demonstrate specific expression in any part of the developing teeth [65], however the gene *WIF1 (WNT* Inhibitory Factor 1) is relatively close by and has a bivalent chromatin status we previously showed to be enriched for craniofacial relevant genes [40]. Notably *Wif1* has been shown in mice to be expressed strongly in the enamel knot, an non-proliferative signaling center required for normal tooth development, where it plays a role in normal tooth morphology (Figure 1C)[71]. This variant is within strong linkage disequilibrium of many other odontogenesis-associated variants which overlap human craniofacial enhancers that are components of larger human craniofacial superenhancer regions. The mouse ortholog of this locus also demonstrated craniofacial enhancer activity, and the larger chromatin landscape surrounding these conserved enhancers includes VISTA-validated enhancers with activity in the pre-dental regions of the pharyngeal arches (Figure 1C). These findings suggest that rs12229918 may disrupt a regulatory region interacting with *WIF1*, demonstrating a locus at which craniofacial enhancers and their target genes may play a role in dental malformation. A similar scenario was observed for the strongest caries associated variant which is in strong disequilibrium with a conserved craniofacial enhancer demonstrated to target *Pitx1* in mouse (Supplemental Figure 2).

### Section 2. Tooth-specific enhancers and disease burden

While these findings were encouraging, they were identified in bulk mouse and human craniofacial tissue up to and including the approximated emergence of dedicated dental cell types. We hypothesized that the enrichment of DVs would be even more pronounced in isolated tooth tissue, particularly tooth-specific enhancers. While it would be ideal to perform this analysis on human developing teeth, the scarcity of this tissue necessitates the use of an animal model. As such, we relied on the mouse model, which is the standard in the tooth development field.

To identify enhancers active in the developing tooth at early stages, we isolated and pooled E13.5 mouse mandibular incisors and performed chromatin immunoprecipitation and sequencing (ChIP-seq) paired with a standard data analysis pipeline (Figure 2A)(Methods) [40,52,53]. We identified 25,358 strong tooth enhancers (TEs), including 6236 enhancers which were unique to the developing tooth compared to hundreds of other samples (tooth-specific enhancers, TSEs; Methods)(Supplemental Table 2, 3). To confirm we had identified tooth enhancers, we interrogated the genome-wide overlap of TEs for validated VISTA enhancers [66] which demonstrated positive enhancer activity in the craniofacial region. We observed a significant enrichment of craniofacial-positive VISTA enhancers in TEs (p<2.2e-16; odds ratio=3.55, Fisher Exact Test). Notably, we observed the same phenomenon in TSEs, despite the significantly decreased number of enhancers. These TSEs were even more highly enriched for craniofacial-positive VISTA enhancers (p=2.391e-09; odds ratio=4.71, Fisher Exact Test)(Figure 2B, Supplemental Table 4, https://cotney.research.uchc.edu/tooth/). An interactive table of these VISTA-validated elements overlapping TSEs and their images from VISTA can be explored at our website (https://cotney.research.uchc.edu/tooth/).

**Figure 2.**
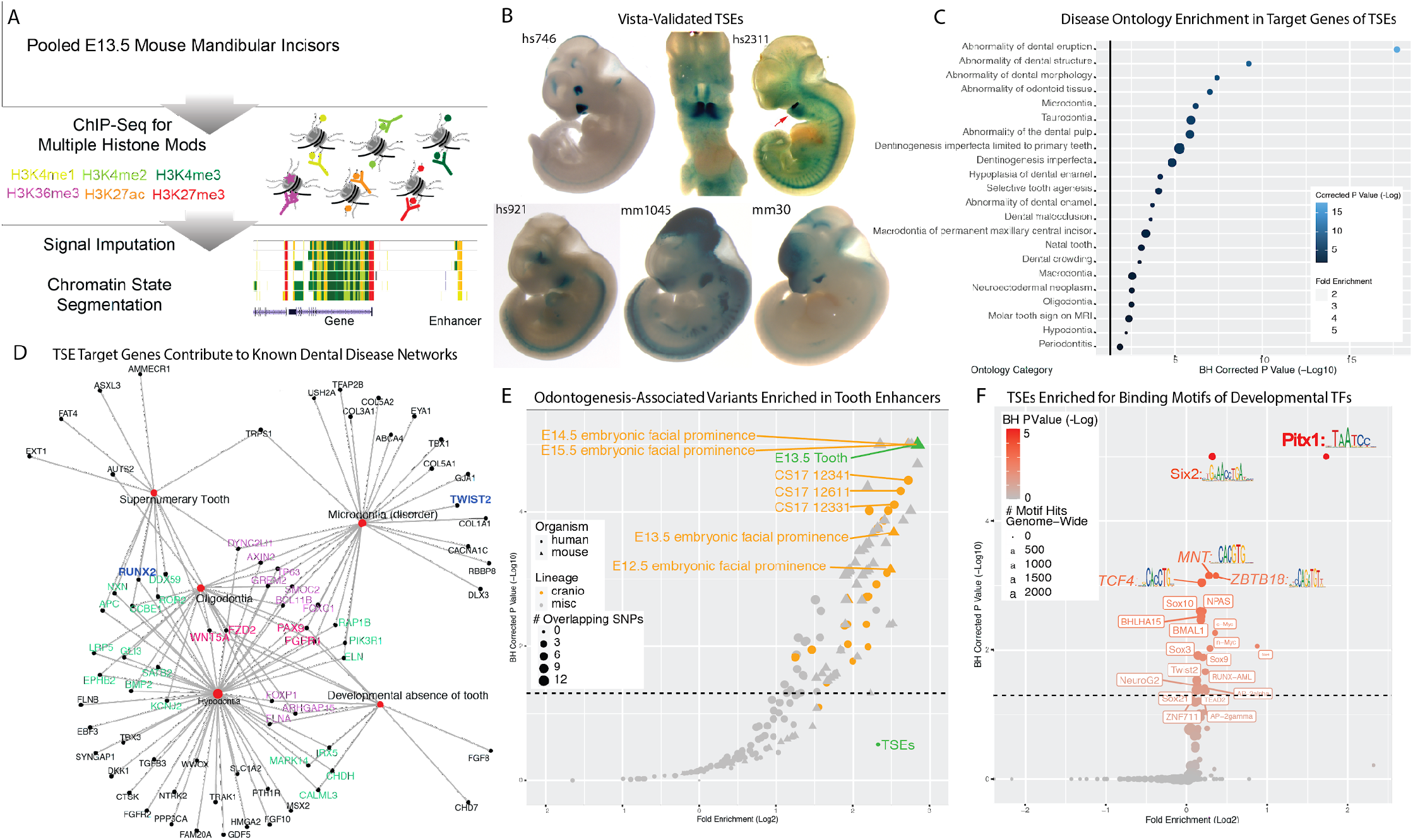
Enhancers of the Developing Tooth Are Enriched For Dental Phenotype Variants and Target Tooth Development Genes A. Schematic of ChIP-seq experiment and chromatin state annotation pipeline of E13.5 mouse mandibular incisors. B. Selected VISTA-validated [66,72] craniofacial enhancers which overlap TSEs. C. Dental disease ontology categories enriched in putative target genes of TSEs, as assigned by Genomic Regions Enrichment of Annotations Tool (GREAT) [72]. D. Detailed gene networks of putative TSE target genes contributing to dental disease ontology categories from C. E. GREGOR [64] analysis of enrichment of “Odontogenesis” (GO_0042476)-associated variants from GWAS Catalog in active enhancers annotated by 18 state chromatin segmentations genome-wide for samples from [40] (orange circles), Roadmap Epigenome (grey circles), and our own 18 state segmentations from ENCODE craniofacial ChIP-seq samples (orange triangles), E13.5 incisor (green) and other samples (grey triangles). Tooth-specific enhancers (TSEs, green) do not pass the significance threshold of 1.3 (dotted line). F. Hypergeometric Optimization of Motif Enrichment (HOMER) [73] analysis of known transcription factor binding motifs enriched within tooth specific enhancers. Position weight matrices are demonstrated for each motif, obtained from JASPAR [74].

Previous publications have shown the enrichment of tissue-specific enhancers around genes with high tissue importance [53,70,75]. Therefore, we hypothesized that the genes targeted by tooth-specific enhancers would be enriched for genes important to dental development and disease. We used GREAT [72](Methods) to assign putative target genes of TSEs (Supplemental Table 5). These putative gene targets were enriched for tooth-relevant gene and disease ontologies, including known dental developmental diseases such as taurodontia, tooth agenesis, and abnormality of the enamel and dentin (Figure 2C,D; Supplemental Table 6)(Methods).

With these positive results of the validity of TEs and TSEs, we next aimed to test the enrichment of DVs within mouse tooth enhancers. To achieve this, we obtained the equivalent hg19 coordinates of tooth enhancers with at least 25% sequence conservation (Methods), resulting in 22,001 conserved TEs and 5,458 conserved TSEs (Supplemental Table 7, 8). We then interrogated these regions for enrichment of DVs using the same approach described above. We observed the highest enrichment (7.17 fold enrichment; padj<0.05) of odontogenesis-associated variants within conserved TEs (Figure 2E). When we re-examined the decreased tooth number-associated locus surrounding *WIF1* (Figure 1C) for conserved TSEs, we noted a high number of these regions predicted to target *WIF1* (Supplemental Figure 3). These results further suggest the validity of the findings and support the conservation of the role of dental developmental enhancers in dental phenotypes.

Enhancers which are active in one specific tissue compared to many others often demonstrate enrichment of binding motifs for transcription factors which are differentially expressed in the same tissue [53]. Therefore we hypothesized that TSEs would be enriched for binding motifs of transcription factors with known roles in dental development and disease. As such, we interrogated enriched transcription factor binding motifs in all TSEs genome-wide using Hypergeometric Optimization of Motif Enrichment (HOMER)[73]. As expected, TSEs displayed enrichment of binding motifs for transcription factor families *Pitx, Sox, Tead, Six*, (Supplemental Table 9, Figure 2F)(Methods). These TFs are relevant to known dental developmental pathways [69,76,77]. The presence of many of these genes both as TSE targets and enriched binding motifs in TSEs reinforce their relevance in the developing tooth structure.

Together, these findings confirmed the role of developmental tooth enhancers in risk of dental phenotypes. These findings additionally suggest the developing mouse tooth is an appropriate model system for the role of enhancers in human dental phenotypes.

### Section 3. WGCNA Reveals Tooth-Relevant Co-Expressed Gene Modules

These findings indicated that developmental tooth enhancers target canonical dental developmental genes. The human orthologs of many of these genes have known dental implications, suggesting TSEs likely contribute to dental phenotypes through target gene dysregulation. As previously mentioned, currently known dental phenotype-driving genes are generally associated with severe syndromes. Therefore we looked to identify previously undescribed genes which may contribute to common dental phenotypes. For genes with unannotated roles in dental development, we looked to integrate our ChIP data with transcriptomic data, specifically using weighted gene co-expression network analysis (WGCNA, Methods)[78]. Gene coexpression analysis performed on samples spanning the development of an organ have been shown to identify both known and putative novel phenotype-relevant genes, as well as cell type specific genes which tend to be coexpressed over time [53]. This has previously been applied in the developing brain and heart, where it has successfully identified genes which likely contribute to autism spectrum disorder and congenital heart disorders, respectively [53,79–81].

For WGCNA, we obtained publicly available bulk RNA-seq data from timepoints spanning the majority of dental developmental stages from both mandibular incisors and molars [82,83](Methods), including rudimentary and relatively differentiated timepoints (E12.5 and E17.5, respectively)(Figure 3A). We aligned, filtered, corrected batch effects, normalized counts, and applied principal component analysis (PCA) to identify major sources of variation in the data (Methods). The resulting PCA demonstrated the corrected batch effect (Supplemental Figure 4). We then applied WGCNA (Methods), which resulted in 28 unique modules of coexpressed genes, spanning 22016 expressed genes (Supplemental Table 10). These modules range from sizes of 63-5330 genes with a median 361 genes/module. To determine their biological relevance, we interrogated the enrichment of gene ontology categories for all genes of each module (Methods)(Supplemental Table 11). 19/28 modules demonstrated enrichment of at least one GO term (padj < 0.1, Figure 3A) relevant to biological processes in the developing tooth. Many of these processes are known to be contemporaneous and interdependent in the tooth, for example epithelial development (midnight blue) and skeletal development (red), which is reflected in the inter-module correlation of Figure 3B. These results highlight the dental significance of a subset of WGCNA coexpression modules, which is demonstrated for specific dental diseases in Figure 3D.

**Figure 3.**
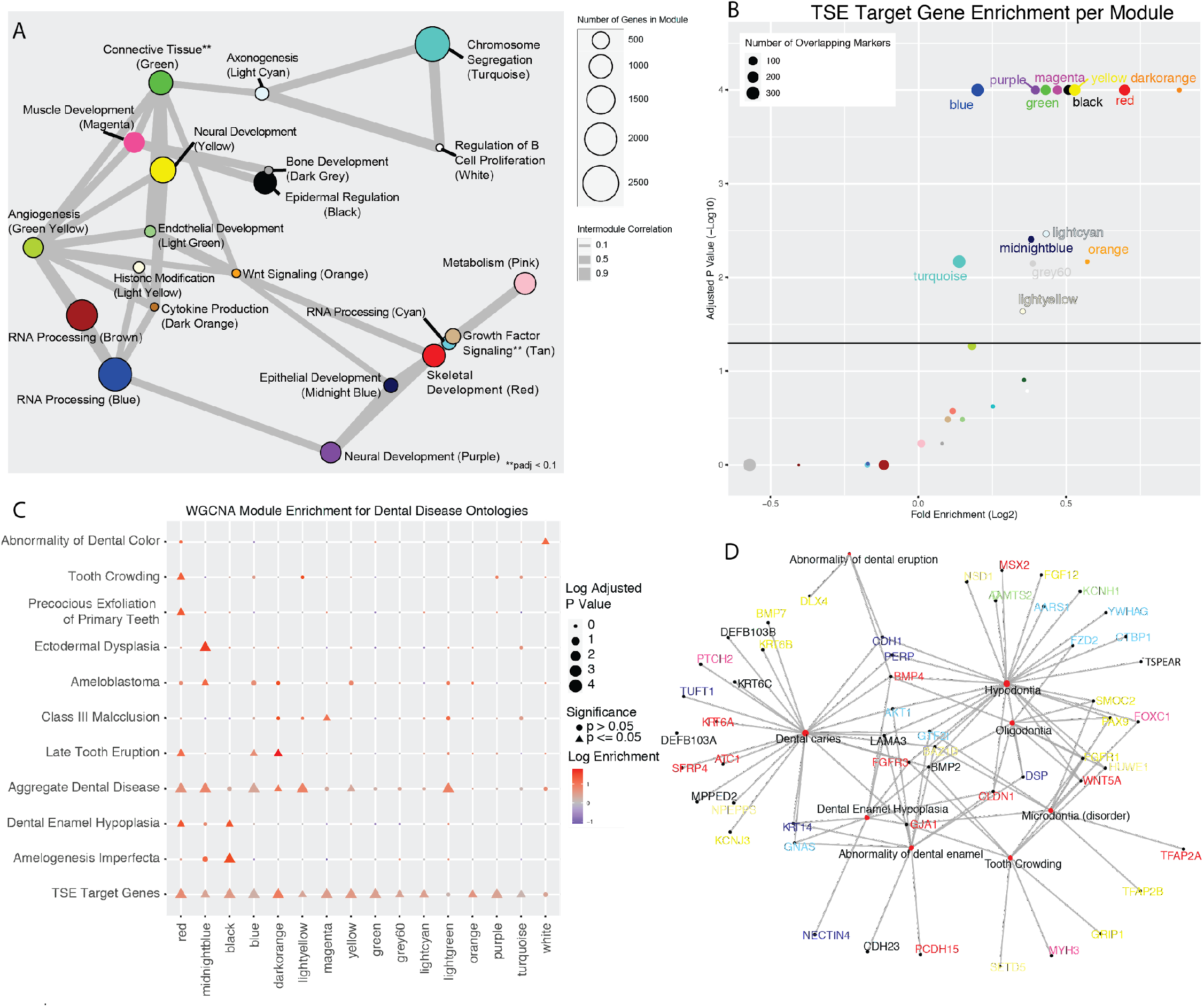
Weighted Gene Co-Expression Network Analysis Reveals Dental Development-Specific CoExpression Modules A. Network plot of co-expressed modules of genes identified through WGCNA on embryonic mouse mandibular tooth samples from molars (E12.5-E17.5) and incisors (E12.5). A Pearson correlation of the module eigenvectors was calculated for the edges, and positive correlations of ≥0.5 were included. The location of each module is determined by multidimensional scaling (MDS) of the module eigengene vectors. Size of dots indicates the number of genes in each module, and colors are module names assigned by WGCNA. Modules are labeled according to module name and the most representative significantly enriched GO term of that module, as determined by clusterProfiler. B. Permutation enrichment analysis (n=10,000) of genes from each module within TSE target genes. Points are named according to the WGCNA module, and point size indicates the number of genes per module which appear as a TSE target gene. C. Permutation enrichment analysis of orthologs of known human dental disease gene categories (Methods) across WGCNA modules. Point size and shape indicates the significance of this enrichment, and color indicates the directionality of that enrichment (positive or negative). D. Detailed gene networks of genes from WGCNA modules in C contributing to human dental diseases. Genes are colored according to their module assignment.

Given the relevant GO enrichments, we looked to determine which modules contained genes specifically active in tooth development. To answer this question, we looked at the enrichment of genes at a per-WGCNA module level within the TSE target genes using a permutation test (n=10,000; Methods). As seen in Figure 3C, we observed significant enrichment of genes from 13 WGCNA modules within TSE target genes (Supplemental Table 12). Notably, these significantly TSE-target enriched modules demonstrated significant enrichment of GO terms relevant to tooth development (Figure 3B). These terms included skeletal development (red), epithelial development (midnight blue), epidermal regulation (black), neural development (purple and yellow), and *Wnt* signaling (orange).

While these results suggested the biological relevance of this subset of modules in tooth phenotypes, these GO terms did not directly correlate with dentally relevant human disease findings. To directly determine if these modules were enriched for genes with dentally relevant phenotypes in humans, we interrogated all modules for enrichment of genes implicated in all diseases with dental phenotypes in DisGeNET [84] (Methods). These genes were interrogated both individually and as an aggregate gene list (Supplemental Table 13). As seen in Figure 3C, the majority of these modules (8/13) identified above were enriched for genes implicated in at least one known dental phenotype (Supplemental Table 14). The GO terms enriched in these modules (regulation of b cell proliferation and endothelial development, respectively) are interesting in that they do not necessarily indicate any dental-specific biological processes, but these combined results rather imply that these processes are necessary for phenotypically normal dental development.

The implication of multiple WGCNA modules in dental development through DO, GO, and TSE target gene enrichment indicates that this technique serves to highlight genes which are likely important to the tissue. Their implication in known human dental phenotypes and combinatorial expression over time reflects known biology of a subset of the genes within each module, helping to prioritize modules for further investigation. Specifically, the WGCNA modules implicated by known genes relevant to dental development in mice (GO and TSE target genes) and dental phenotypes in humans (DO) are suspected to include novel genes which may be considered candidate dental disease genes.

### Section 4. scRNA suggests cell type specific contribution of co-expressed gene modules

These combinatorial results of TSE target genes and tooth-relevant WGCNA modules suggest lists of co-expressed genes which are likely important to tooth development and disease. However, the bulk nature of WGCNA obscures the contributing cell types of these important genes. The developing tooth is composed of layers of interdependent epithelial and neural crest-derived mesenchymal cell types, and knowing contributing cell types of candidate disease genes is crucial to developing proper treatment of maldevelopment. The minute size of the structures and this intricacy of the composing cell types necessitated single-cell level transcriptomic analysis to identify the cell types contributing to disease risk.

We reprocessed published scRNA-seq data from four E14 mouse molars [82](Methods). A graph-based clustering approach [85] revealed 33,886 cells separating into 16 transcriptionally distinct cell states (Supplemental Figure 5). Using pseudobulk methods [86] we identified 4821 total marker genes, an average 301 per cluster (Methods; Supplemental Table 15). We leveraged significant GO and DO terms enriched in each cluster (Methods) to assign biological identities per cluster (Supplemental Table 16, Figure 4A,B), and confirmed assignments with canonical marker genes (Supplemental Figure 6) (Methods). To include only cell types with consistent inclusion across all clusters, we excluded the variable “Bone Progenitor Cells’’ cluster (cluster 10), which indicated normal variation in mandibular tissue inclusion during dissection. Given the cap stage of tooth development of these samples we hypothesized that the enamel knot, a signaling center made up of a small set of aproliferative epithelial cells required for normal tooth development, would be present and identifiable within a subset of the epithelium. Using a subclustering approach (Methods), we were able to identify an epithelial subcluster with attributes consistent with canonical markers of the primary enamel knot (Figure 4C, Supplemental Figure 7), which we called the putative enamel knot (PEK). This subset of epithelial cells was therefore included in the annotated UMAP (Figure 4A).

**Figure 4.**
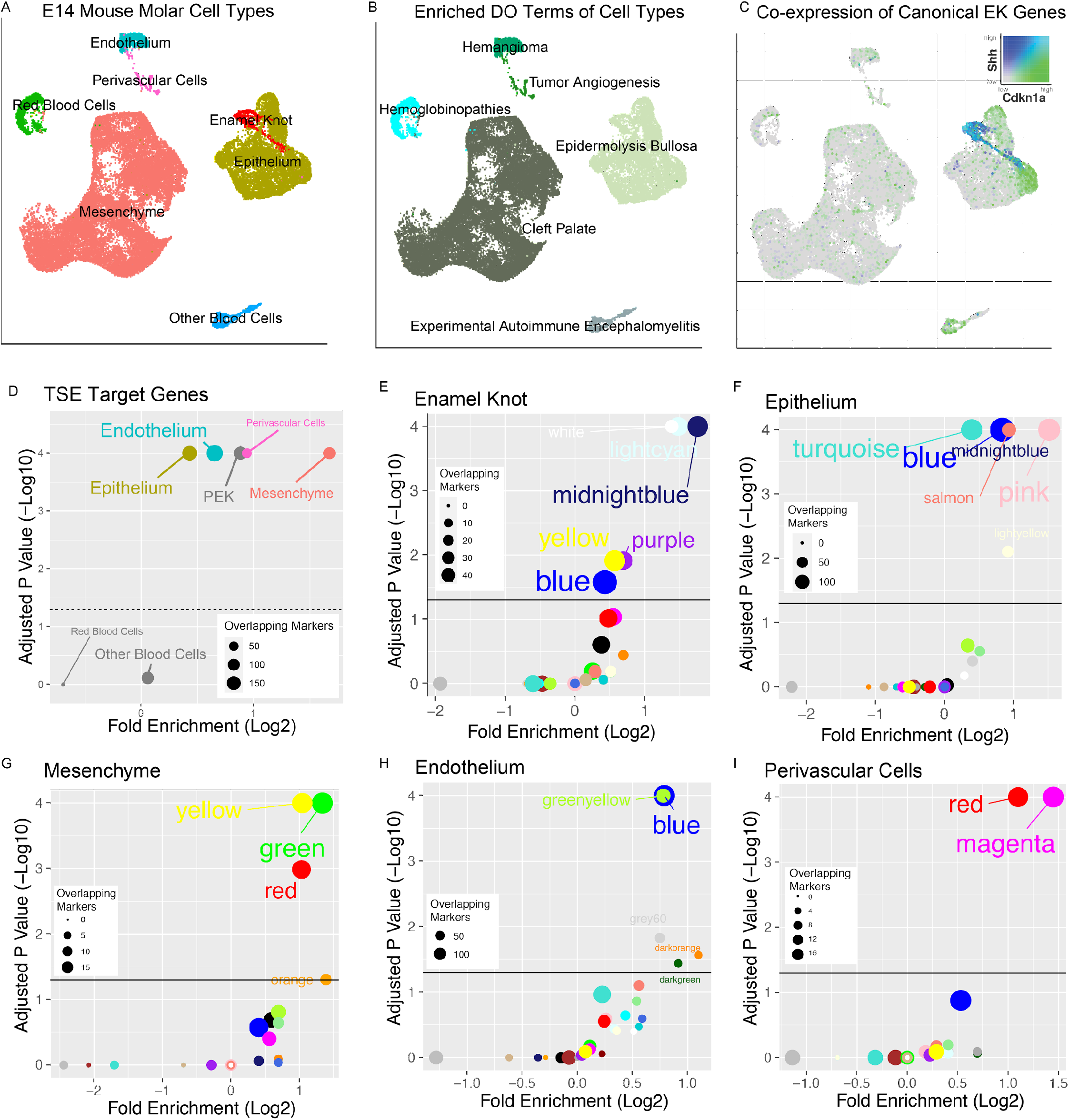
Single Cell Analyses Suggest Cell-Type Specific Activity of WGCNA Modules A. Annotated UMAP of cell types determined through Seurat analysis of 4 replicates of E14 mouse mandibular molars (n=33,886 cells) from [82]. B. UMAP as in A, labeled according to the most representative significantly enriched human disease ontology in each cell type. C. Co-Expression overlay of UMAP from A demonstrates a band of *Shh/Cdkn1a+* (cerulean) epithelial cells that make up the putative enamel knot, consistent with canonical traits of the structure. D. Permutation enrichment analysis of marker genes from each cluster within TSE target genes. Points are named according to the cluster, and point size indicates the number of genes per cluster which appear as a TSE target gene. This indicates that blood cells are not a major contributor of tooth-specific enhancers in bulk analyses. E-I Permutation enrichment analysis of marker genes from each cluster within genes from each WGCNA module. Each plot indicates the major cell type being interrogated. Points are named according to the WGCNA module, and point size indicates the number of genes per module which appear as marker genes in that tissue.

Having annotated the cell types of the E14 mouse molar including the previously un-annotated PEK, we generated transcriptomic profiles of all cell types. We identified 2573 total marker genes, an average of 367 per cluster, including the 317 PEK genes (Supplemental Table 17). As expected, 1:1 human orthologs of these marker genes were enriched for cell type-relevant disease ontologies (Figure 4B). Notably, these marker genes contained canonical EK marker genes, including the previously mentioned *Wif1* (Figure 1C), validating our annotation. As the EK has not previously been profiled in single cell analyses and the vast majority (287/317) of PEK marker genes were novel, we looked to confirm the expression of the top PEK genes (TPEKs; Methods). The 60 TPEKs included 8 canonical EK genes and 52 genes not previously identified in the EK. We performed a literature survey and referenced the GenePaint resource of in situ hybridization (ISH) images of the E14.5 mouse embryo [65,87–90], looking at the approximate area of the mandibular first molar. 47 novel TPEKs had previously been profiled via ISH, the overwhelming majority of which (38/47) showed strong and specific hybridization closely mirroring the results for *Shh, Wif1*, or *Wnt10b* (Supplemental Table 18, https://cotney.research.uchc.edu/tooth/). An interactive table of all TPEKs can be further explored on our website, https://cotney.research.uchc.edu/tooth/.

Having generated transcriptomic profiles of all cell types, we looked to ascertain if the previously identified TSE target genes are enriched for marker genes of any specific cell type. Using a random permutation test (n=10,000, Methods), we revealed a significant enrichment of marker genes from all major, dentally relevant cell types within TSE target genes (Figure 4D)(Supplemental Table 19). Specifically, the only cell types whose marker genes were not enriched within TSE target genes were red blood cells and other blood cells.

With these promising results indicating validity of the identified marker genes, we asked if the prioritized WGCNA modules were enriched for marker genes of any specific cell type. As expected, we observed marker genes to be enriched in WGCNA modules with GO terms relevant to that cell type (Supplemental Table 20, Supplemental Figure 8). For instance, genes of the midnight blue module were enriched for epithelial GO terms and marker genes of both the epithelium and the epithelial-derived PEK (Figure 4D,E). Similarly, genes of the green-yellow module were enriched for angiogenesis-related terms, and marker genes of the endothelial cluster (Figure 4F). This continued with enrichment of the red module (bone GO terms) in the mesenchyme, salmon (cell migration-related GO terms) in other blood cells, and magenta (muscle GO terms) in the perivascular cells (Figure 4G-I). As expected, many of these top enriched modules for the dental-specific cell types were seen to be enriched for TSE target genes (Figure 3). Altogether, these findings suggest that prioritized modules of co-expressed genes are expressed in cell type specific patterns in the developing tooth, where they are likely important to normal dental development.

### Section 5. Data integration suggests novel dental phenotype genes

These layers of transcriptomic data suggested a subset of coexpressed modules of genes for further analysis. However, these modules combined included thousands of genes which required prioritization to inform researchers on promising candidates to pursue. As such, we sought to integrate the multiple presented data modalities (transcriptome, single cell transcriptome, GWAS, ChIP-seq) to generate a candidate disease list. Genes passing a minimum expression threshold in the bulk data (Methods) were prioritized using the following general categories: 1. Classification of the gene as a marker gene from scRNA analysis, 2. Dental relevance of the WGCNA module assigned to each gene, 3. Proximity of each gene’s human ortholog to a craniofacial-relevant variant from GWAS catalog 4. Proximity of each gene to a VISTA-validated TE, and 5. Number of TSEs predicted to target each gene (Methods). Overall, this method combined 5 categories to determine the priority of 16322 genes genome-wide.

The top priority gene through this prioritization method was revealed to be *Runx2*, a canonical dental development gene whose human ortholog is associated with cleidocranial dysplasia, a disease with prominent dental abnormalities [91]. The top 20 genes also included known dental disease genes *Bmp4, Zfhx3, Tbx3, Twist1, Msx2*, and *Postn* which cause phenotypes such as abnormal tooth morphology, dental caries, and tooth agenesis [92–95]. These findings within our top scoring genes validated our prioritization system. We present in Supplemental Table 21 our prioritized list of 1632 top scoring candidate disease genes, including 1416 novel candidate disease genes.

With these promising results, we sought to validate a novel putative disease gene *in silico*. We looked to validate a gene which: 1. Had successfully been knocked out body-wide by the mouse knockout project (KOMP) without significant early lethality [96], 2. Had successfully been assayed by GenePaint in the E14.5 mouse, and 3. Had a VISTA-validated TE within 1Mb of the TSS. We focused upon *Agap1* (Arf1 GTPase activating protein 1), the highest priority gene meeting these criteria.

*AGAP1* in humans contains the top scoring sequence of human acceleration on a conserved background in the genome, called HACNS1 (chr2:236773456-236774696) [97]. This sequence is located in an intron of the gene, and actually lies within a human craniofacial superenhancer region (Figure 5A), suggesting its role in the craniofacial apparatus. Despite its proximity to known craniofacial regulatory regions, HACNS1 has been proposed to regulate *Gbx2* in the mouse limb [98]. However, this gene is expressed at relatively low levels in human craniofacial tissues [70] and the mouse molar compared to *Agap1* (Supplemental Figure 9). These findings suggest that *AGAP1* could play an important role in facial development and morphology and potentially be regulated in a human-specific fashion during tooth development. While *Agap1* does not appear in a module that is enriched in TSE target genes (brown module, Figure 3B), the 1Mb region surrounding the gene contains 7 TSEs which likely target *Agap1*, as it is the only tooth-specific gene in the region (Figure 5A). It is notable that while variants within *AGAP1* itself have not been implicated in human disease, a variant intronic to *AGAP1* has been associated with hemifacial microsomia (rs3754648, p=5e-13), a disease which affects the patterning of the face and often coincides with dental phenotypes [99]. This variant lies within a human craniofacial superenhancer region, and is within linkage disequilibrium of a conserved tooth-specific enhancer that was validated by VISTA (Figure 5A,B) and demonstrates highly tooth-specific activation.

**Figure 5.**
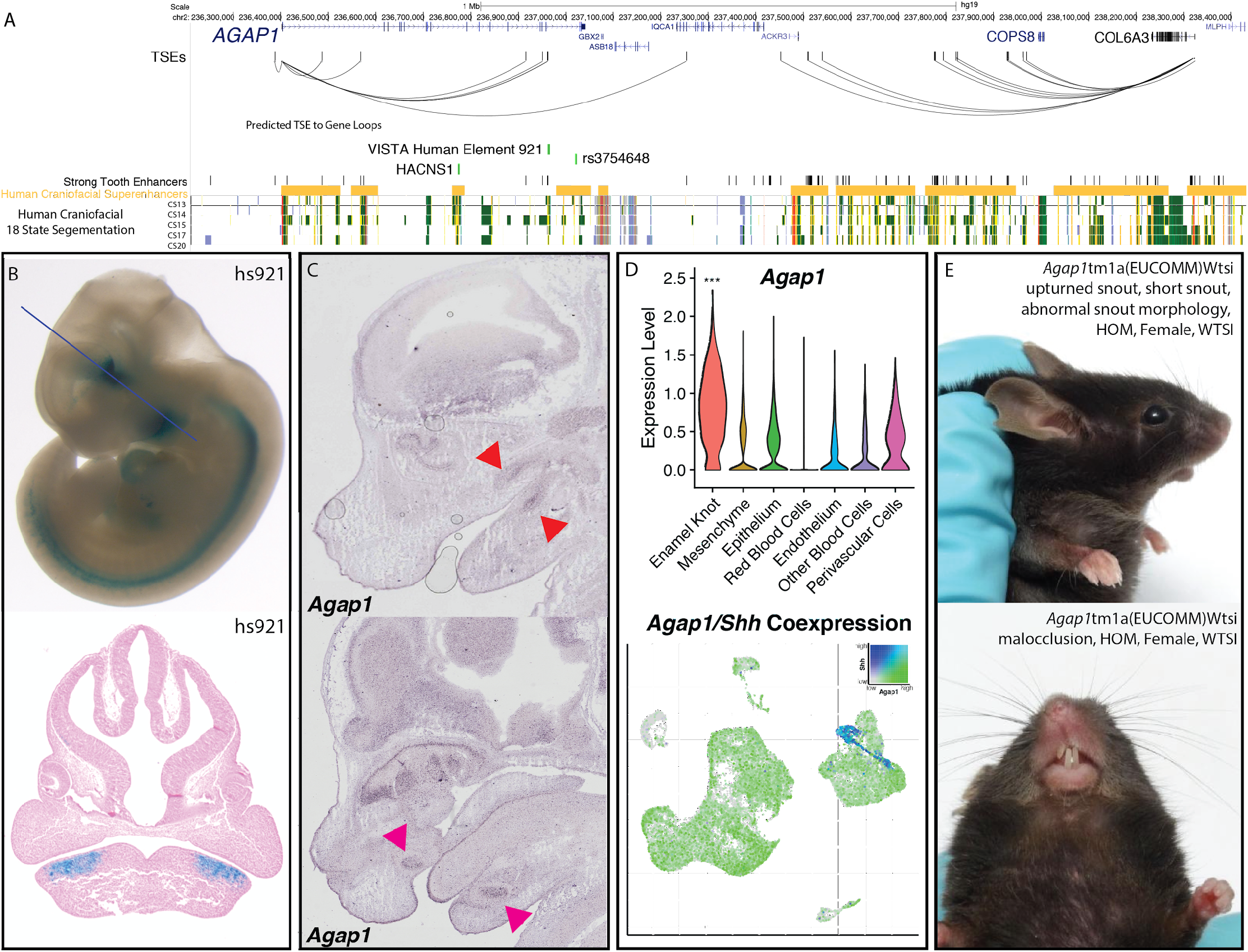
Validation of *Agap1* as a Novel Dental Disease Gene A. Visualization of the hg19 locus surrounding *AGAP1*. TSEs, strong tooth enhancers, and predicted TSE to gene loops were determined as described (Methods) using mm10 coordinates and lifted to hg19. B. VISTA-validated enhancer hs921 demonstrates activity in the pre-mandibular pharyngeal arches, where it is limited to the approximate pre-dental regions. Images from VISTA [66]. C. *Agap1* expression is apparent in both the maxillary and mandibular molars (top, red arrows) and incisors (bottom, pink arrows). *in situ* hybridization images from GenePaint [65,87–89]. D. Single cell transcriptomic analysis demonstrated the PEK-specific expression of *Agap1* in the E14 mouse molar. p<0.05, Wilcoxon test. E. *Agap1* loss of function mutations in mice result in malocclusion and abnormal facial feature phenotypes at 13 weeks postnatal age. Images from KOMP [96].

Despite the presence of a relevant variant within the gene, *AGAP1* has no reported dental phenotype in humans, and has not been associated with any specific human syndrome. Interestingly, the gnomAD entry for *AGAP1* demonstrates a LOEUF score (0.23) [100] in the bottom decile of all genes, indicating the gene is resistant to loss of function mutations and therefore likely extremely important in humans. *Agap1* has also been demonstrated to be expressed in the upper and lower incisors and molars at E14.5 [65,87–90], where it appears to be restricted to the previously described approximate enamel knot area (Figure 5C-D, Supplemental Figure 10). Supporting this finding is its high expression in the PEK; *Agap1* appeared in our single cell analysis as a PEK marker gene, with 0.7 log fold enrichment in the PEK compared to all other cell types (padj=1.51e-70)(Figure 5D). Many genes specific to the EK result in dental phenotypes when knocked out specifically in the tooth [101,102], and the specificity of this gene to the PEK within our data suggests there would likely be a similar phenotype.

To determine if this is the case, we turned to the mouse knockout project (KOMP) to discern whether dental related phenotypes were apparent upon disruption of *Agap1*. Exploration of the database entry for the homozygous loss of function *Agap1* mice revealed clear and well described craniofacial phenotypes of snout malformation and malocclusion (improper meeting of maxillary and mandibular teeth) (Figure 5E, Supplemental Figure 11). Results from the KOMP also cited increased prevalence of “abnormal tooth morphology” (p=9.12e5).

Altogether, these factors in combination suggest *AGAP1* is an important player in dental development in humans and mice, and that modulating its expression likely results in a dental phenotype. More importantly, *AGAP1* serves as a validation of our results from the integration of multiple datasets from mouse tissue. These results also indicate the parallels this data is able to draw in the congruent human tissue when that tissue is unable to be obtained.

## Discussion

Previous findings in the bulk developing human craniofacial regions demonstrated a role for active regulatory regions in craniofacial morphology and disease. Here we have shown that these regions likely also influence dental phenotypes and disease. Using publicly available ChIP-seq data we have generated functional chromatin annotations in hundreds of mouse tissues including craniofacial samples. With these annotations we have demonstrated that conserved regulatory regions active in the early developing mouse face are similarly systematically enriched for variants associated with a variety of human dental phenotypes.

We performed ChIP-seq on isolated E13.5 mouse incisors in order to understand the role of the enhancers from the tooth itself in determining these dental phenotypes and diseases. We observed that this tissue demonstrates the highest enrichment of odontogenesis-associated variants, suggesting dysregulation of enhancers specifically active in the tooth contributes to these tooth-isolated phenotypes. This conservation of findings across species indicates that the mouse tooth is a viable model for the role of enhancers in dental development and disease. To determine the genes which are likely contributing to these phenotypes, we leveraged publicly available bulk and single cell transcriptomic data, performed gene co-expression analysis and identified cell type specific transcriptomic signatures. We integrated these findings with our tooth enhancers and previously generated GWAS for dental phenotypes to prioritize genes based on their likelihood to be relevant to tooth development. While data integration of this type has proven to be useful in identifying human disease candidate genes from developing tissue [53,70], this method has not been applied to the developing mouse tooth, nor has it been extrapolated to indicate human dental disease genes.

We have demonstrated here that variants associated with odontogenic phenotypes are enriched in enhancers active in the human and mouse developing craniofacial apparatus. This role for enhancers in tooth phenotypes has not previously been demonstrated, and the role of enhancers in tooth development as a whole has been largely unexplored. In this manuscript we described a specific locus which serves as an example of the role these regions can play in dental phenotypes. We observed a variant highly associated with decreased tooth number upstream of *WIF1*, a gene known to be isolated to the enamel knot in mice where it plays an important role in tooth morphogenesis (Figure 1C). This variant and others within high linkage disequilibrium were found to overlap with conserved regions of craniofacial enhancers, many of which were predicted to target *WIF1* (Figure 1C, Supplemental Figure 3). Interestingly, this gene has not been described in human dental development or disease. After integration of all our data modalities in this investigation, this gene appeared in the top ten of our candidate disease gene list, further suggesting this gene’s role in dental development and phenotypes is conserved across species.

We have also shown here that active enhancers regulating patterning of the human face also harbor a significant number of variants related to caries risk, a disease whose risk is generally considered to be both developmental and immune-mediated [13,103–105]. However, the majority of cases do not demonstrate a single-gene etiology. Our findings suggest that craniofacial enhancers play an important role in this risk, supported by our specific findings at the *PITX1* locus (Supplemental Figure 2). The location of the strongest caries-associated variant in a conserved craniofacial enhancer region that has been shown to influence *Pitx1*, an epithelial marker gene. This suggests that modulation of its expression affects the development of dental epithelial cell types in a way which may contribute to risk of caries over time. Further, it is important to note that we observed enrichment of caries-associated variants in immune cell enhancers in addition to craniofacial enhancers (Figure 1B). Given that we observed no enrichment of TSE target genes within markers of red blood cells or immune cells (Figure 4D), we determined that these cell types do not largely contribute to enhancers identified in the bulk tooth. Rather, our results suggest a duality of independent genetic caries risk contribution from the developing tooth (prenatal risk) and immune system (postnatal risk). Additionally, we provide a list of novel genes likely important to tooth development which may be of interest for downstream modulation investigations in caries models.

In this manuscript we have also presented a re-analysis of publicly available scRNA-seq data from the E14 mandibular molar, from which we have identified a novel putative enamel knot gene signature. Using canonical EK marker genes, we annotated a previously undiscovered epithelial subcluster with features consistent with the EK (Figure 4C, Supplemental Figure 7). We uncovered 317 PEK-specific genes and demonstrated the relative EK specificity of the majority of the 60 most specific PEK genes by comparing their expression within the approximate mandibular molar to *in situ* patterns of canonical EK markers *Shh, Wnt10a, and Cdkn1a* (Supplemental Table 18). Given the known role of the structure in patterning and morphogenesis of the tooth [106–108], this novel PEK signature suggests targets for future studies in dental regeneration and morphology, and is available on our website as an interactive table for further exploration.

Lastly, we have here demonstrated the ability to extrapolate multimodal data from the developing mouse tooth to human dental development and malformations. Previous investigations in human tooth development have been limited due to scarcity of the tissue, but they demonstrated the conserved expression of important dental factors such as RUNX2, WNTs, and BMPs [42–46]. Many of these genes appear as marker genes for the mesenchyme, enamel knot, and epithelium, and appear as highly important genes in our prioritized gene list (Supplemental Table 21). This speaks to both the robust nature of our analyses and indicates the validity of this prioritized list. Additionally, our findings genome-wide and our detailed findings at the *PITX1* and *WIF1* loci suggest the validity of the mouse tooth as a model of enhancers’ role in human dental development and disease. Further downstream experiments and analyses will be necessary to unravel the specific roles of these prioritized candidate disease genes and tooth-specific enhancers in dental development and phenotypes. Additionally the tooth-specific enhancers we have identified could be attractive molecular tools for driving highly specific expression of reporter genes or Cre recombinase as we have shown for precise regions in the mouse brain [54]. We provide a convenient portal to explore all of our results at https://cotney.research.uchc.edu/tooth/ with minimal computational effort for the convenience of the research community.

## Materials and Methods

Detailed scripts of all analyses can be found at our github (https://github.com/cotneylab/tooth).

### Chromatin State Segmentation

E13.5 mouse mandibular incisors were isolated bilaterally from multiple C57BL/6J embryos and crosslinked using 1% formaldehyde as previously described [109,110]. Upon isolation of nuclei and shearing, soluble chromatin was divided equally across multiple tubes for immunoprecipitation with antibodies against H3K4me1, H3K4me2, H3K4me3, H3K27ac, H3K27me3, and H3K9me3. Subsequent sequencing, alignments, imputation of signals, and chromatin state segmentation using 15 and 18 state models were performed as previously described [40]. The same chromatin signals were retrieved from all available data in mouse ENCODE [63] for multiple replicates of 12 tissues and 10 timepoints and imputed to yield 1316 distinct chromatin signals. These imputed signals were then segmented with the same models as above resulting in 154 chromatin state segmentations (https://genome.ucsc.edu/cgi-bin/hgTracks?hgS_doOtherUser=submit&hgS_otherUserName=Jcotney&hgS_otherUserSessionName=Mouse_chromhmm_15_18_state). Tooth specific enhancers were identified using bedtools (v2.29.0, [111]) on strong enhancer annotated segments (18 State chromatin segmentations; States 8, 9, 10; [53]) and compared to other non-craniofacial mouse tissues (samples without any dental tissue) and the same chromatin state segments. Mouse strong enhancers were converted to hg19 coordinates (kent-tools, v1.04.00; minimum sequence conservation 25%) for GWAS analyses (see below). Target genes of these enhancers were identified using rGREAT (v1.26.0, [72,112]) for mm10, using the default settings.

### GWAS enrichment analysis

Caries, Crohn’s disease and odontogenesis-associated variants were obtained from GWAS catalog (EFO_0003819, EFO_0000384, GO_0042476). Variants identified in eastern european cohorts with p <= 5e-8 were retained for each category. GREGOR (v1.4.0, [64]) was used to interrogate enrichment of disease-associated variants within strong enhancers of each biosample, with the control group of Eastern European background variants. Human enrichment analysis was performed on 18 state segmentations of published samples obtained from [40,53] and 127 human cell types profiled by Roadmap Epigenome [52,113]. Mouse enrichment analyses were performed on 18 state segmentations from published mouse tissues from the ENCODE project and our E13.5 mouse incisor sample, as described above.

### WGCNA

Bulk RNA-seq fastqs from molars (n=30; 3-E12.5, 7-E13, 7-E14, 3-E15.5, 7-E16, 3-E17.5) and incisors (n=3, E12.5) from [82,83] were obtained from GEO and aligned using Kallisto (kallisto v0.46.2) to the mm10 genome (GRCm39)[114]. Aligned libraries were imported to DESeq2 (v1.26.0) [115] using tximport (v1.14.2) [116] and one replicate of E13 was omitted as an outlier. Genes were filtered for minimum transcript count of 5 in at least 3 replicates. Batch effects were corrected removing PC1 and PC2 as sources of variation using RUVseq (v1.20.0) [117]. The remaining 32 samples were processed using DESeq2. Variance stabilized transformation count matrix for all samples were imported into a WGCNA pipeline (v.1.70-3) [78]. In brief, samples were filtered for outliers and genes were filtered for quality. Power for each file was obtained and a threshold of 0.85 was used to define power of 8. WGNCA network was built, using unsigned TOM, minimum module size of 50, gene dendrogram merge cut height of 0.25, and a deepSplit of 2. Correlation and intramodule connectivity for each module was identified. Gene ontology enrichment analysis and subsequent figures for each module were generated using clusterProfiler, with “enrichGO” and “enrichDGN” modalities on genes and their 1:1 human orthologs (clusterProfiler v3.14.3, org.Hs.eg.db v3.14.0, org.Mm.eg.db v3.10.0)[118–120].

### Single cell RNA-seq

Raw fastqs were retrieved from GEO (GSE142201) [82] and aligned to the mm10 genome (GRCm39) using Kallisto/bustools (kallisto v0.46.2, bustools v0.40.0)[114], and Kallisto for bulk RNA-seq libraries. Kallisto/bustools was used in filtering and Seurat object generation. Seurat (v3.2.0)[87] was used to merge replicates. Cells were filtered (<5% mtDNA, >200 nFeatureRNA) and log normalized. Cell cycle stages were assigned with mouse orthologs from known human cell cycle genes. Data was scaled using all genes. Nearest neighbors were assigned using dimensions 1:15 via Louvain algorithm. Clusters were generated at resolution=0.3. UMAPs were generated with dimensions 1:15. Epithelial analysis was clustered at resolution=0.2, using dimensions 1:15. Marker genes were identified with DESeq2 using logfc>0.5, min.pct=0.25. SymIDs were converted to EntrezID with clusterProfiler. clusterProfiler was used to calculate GO term enrichment against background. clusterProfiler “simplify” combined similar GO terms, with a z score cutoff of 0.5. Marker genes of annotated cell types were identified after dropping all cells with the identity “Bone Progenitor Cells” from analyses, using DESeq2 with logfc>1.0, min.pct=0.25, max.cells.per.ident=400.

### Marker Gene Validation

In situ hybridization images for EK marker gene validation were obtained from the GenePaint online resource and a literature search for specific genes in the developing tooth [65,87–89][90,121].

### Disease Gene Prioritization

All genes from WGCNA analyses, excepting genes belonging to the grey module (low expression module) were considered for prioritization. Genes were given a “score” based upon a number of factors. Genes were given one point for being a marker gene for any scRNA-seq annotated cell type. An additional point was given for marker genes of dental-specific cell types (Enamel Knot, Epithelium, Mesenchyme, and Perivascular Cells). We obtained hg19 coordinates of GWAS variants associated with craniofacial and dental phenotypes (EFO_0009892, EFO_0600095, GO_0044691, EFO_0009331, Orphanet_141136, and HP_0000175) and identified human genes located within 500kbp of at least one variant (Supplemental Table 22, 23). Mouse orthologs of these genes were given an additional point. Genes were given additional points based on the number of TSEs predicted to be associated by GREAT: 0 TSEs = 0 points, 1-2 TSEs = 1 point, 3-4 TSEs = 2 points, 5-6 TSEs = 3 points, 7+ TSEs = 4 points. Genes belonging to tooth-relevant WGCNA modules (blue, purple, green, magenta, yellow, black, red, midnightblue, darkorange, orange, lightyellow, lightgreen, white, turquoise, lightcyan, grey60) were given an additional point. We used mm10 coordinates of all VISTA+ tooth enhancers to identify all genes within 1Mb of each enhancer (Supplemental Table 24). These genes near VISTA validated enhancers were given a point. These scores were summed for each gene, and genes with the same score were further prioritized based on the predicted number of targeting TSEs.

Statistical significance for enrichment analyses was assigned using random permutation analysis (n=10,000) with Benjamini-Hochberg correction for all tests.

Figures were generated using ggplot2 v3.3.3 and ggsignif v0.6.1. ShinyCell (v.2.0.0)[122] package was used for development of the web portal. Scripts for computational analyses are available at https://github.com/cotneylab/tooth.

## Acknowledgments and Funding

We would like to give our profound thanks to Dr. Mina Mina of UConn School of Dental Medicine for her work in providing the primary embryonic mouse tissues used in these experiments.

We are supported by NIH grants 1R01DE028945, 1R03DE028588, 5R35GM119465, 5T90DE021989, and 1F30DE031149.

---

**Supplemental Figure 1.**
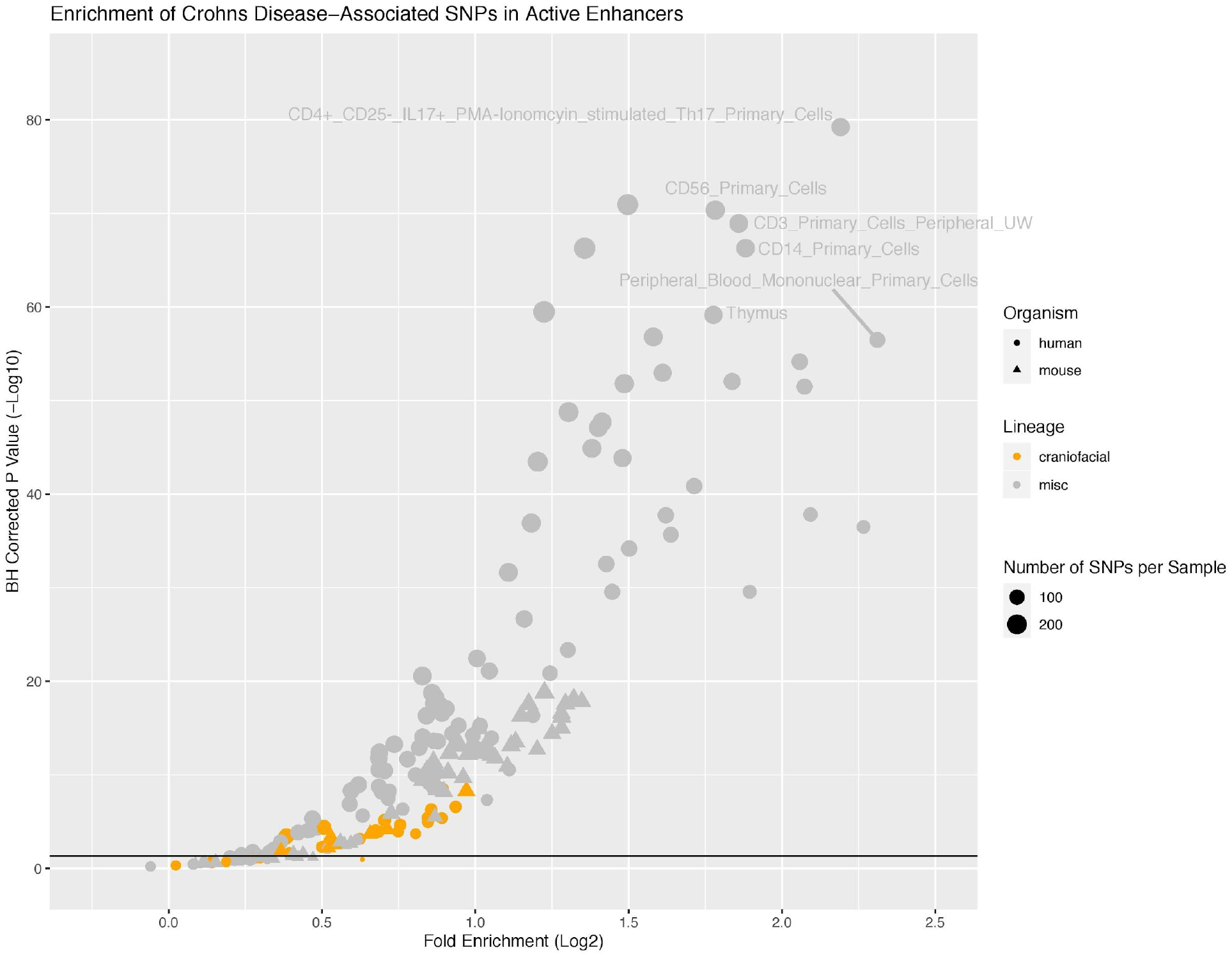
Scatterplot of GREGOR (genomic regulatory elements and GWAS overlap algoRithm) analysis of enrichment of “Crohn’s Disease” (EFO_0000384)-associated variants from GWAS Catalog in active enhancers annotated by 18 state chromatin segmentations genome-wide for samples from [40](orange circles), Roadmap Epigenome (grey circles), and our own 18 state segmentations from Mouse ENCODE craniofacial (orange triangles) and other mouse tissue samples (grey triangles) [40,52,63]

**Supplemental Figure 2.**
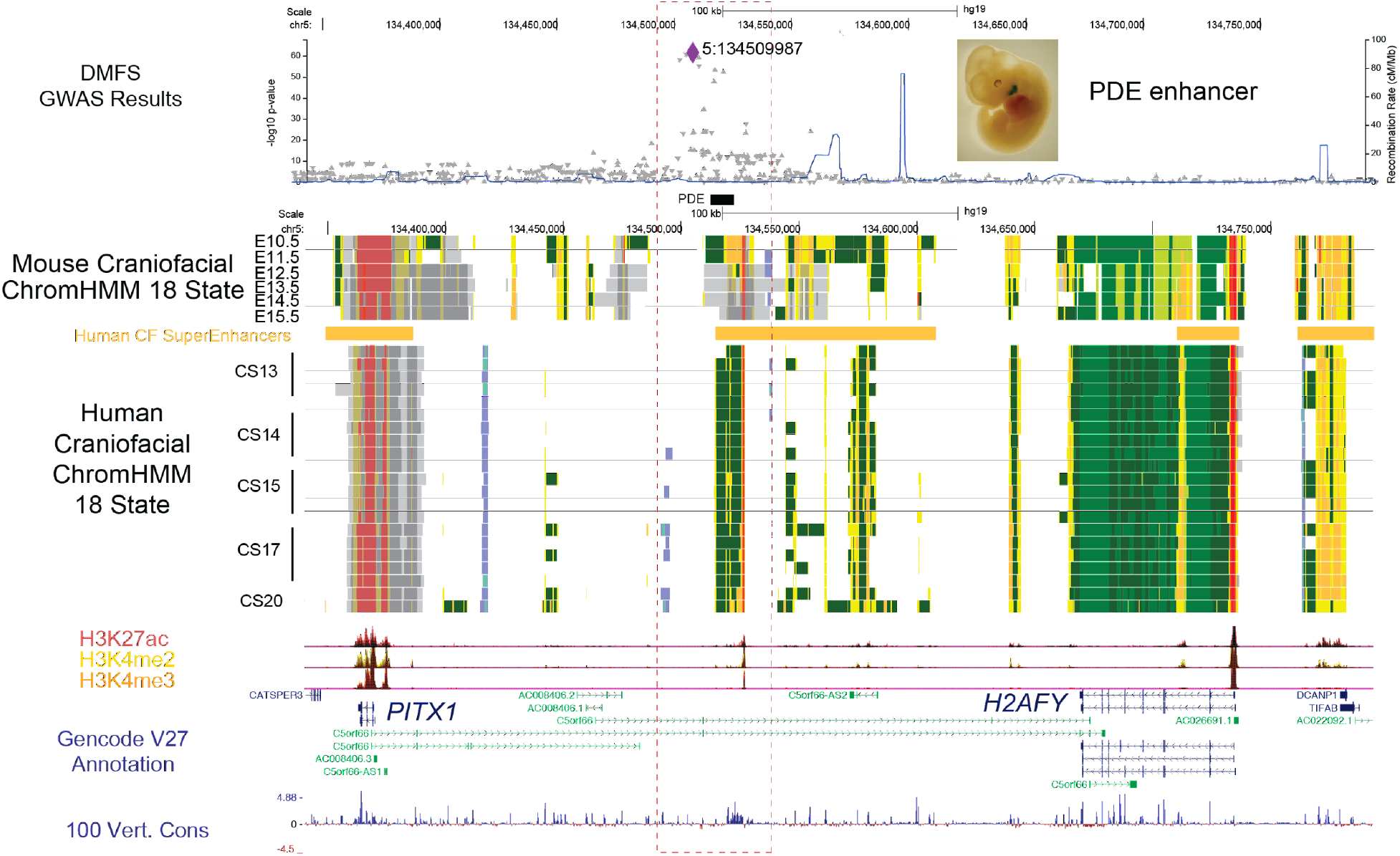
Visualization of the locus surrounding rs1122171 (chr12:65,761,808, hg19) and nearby fine-mapped variants associated with decayed, missing and filled surfaces (DMFS) by [56]. Mouse craniofacial 18 state segmentations were generated as described in Methods from publicly available data from ENCODE [63]. Human craniofacial 18 state segmentation and composite H3K27ac, H3K4me2, and H3K4me3 p-value signal tracks were obtained from [40]. The variant rs1122171 had the strongest association with DMFS and was initially reported to be associated with the uncharacterized gene *C5orf66 [56]*. Upon inspection of our data we found other highly significant DMFS-associated variants in strong linkage disequilibrium with this variant, many of which overlapped a human craniofacial enhancer annotation that is a component of a larger craniofacial superenhancer region [40]. The mouse ortholog of this region has also been shown to have enhancer activity restricted to the developing pharyngeal arches in the very early stages of dental development (E11.5), where it interacts directly with *Pitx1* nearly 200kb away [67](Figure 1C, inset). The region is well conserved across vertebrates and mutations in *PITX1* have similar consequences in humans and mice [68], including roles in tooth and limb development [69]. Additionally, *PITX1* has a bivalent chromatin status in developing human craniofacial tissues and has peak expression at Carnegie stage 17 [70]. These findings suggest that rs1122171 may disrupt a regulatory region interacting with *PITX1*, demonstrating a locus at which craniofacial enhancers and their target genes may play a role in dental malformation and disease.

**Supplemental Figure 3.**
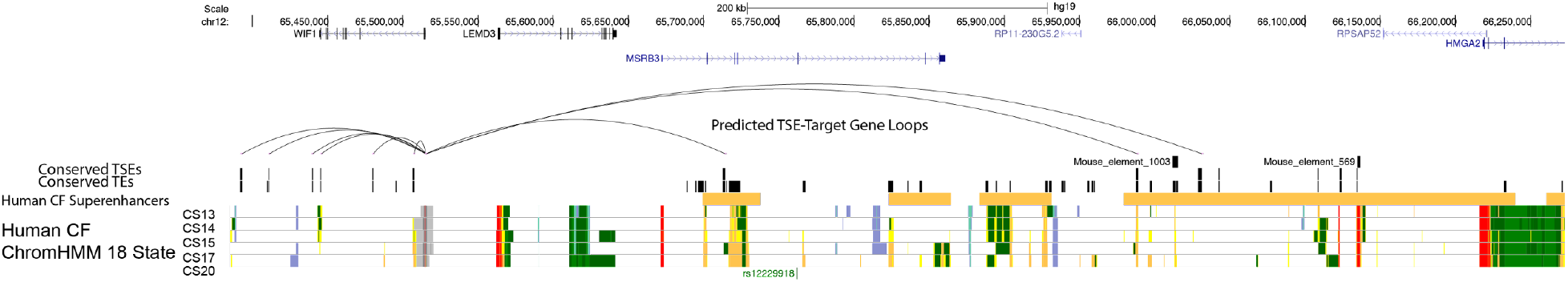
Visualization of the locus surrounding rs12229918 (chr5:134509987, hg19) and nearby fine-mapped variants associated with decreased tooth number and delayed tooth eruption (Odontogenesis) by [57]. Loops demonstrate predicted conserved mouse tooth specific enhancer-target gene interaction. “TEs” track indicates conserved active enhancers of mouse incisor. Human craniofacial superenhancers and 18 state segmentation tracks were obtained from [40].

**Supplemental Figure 4.**
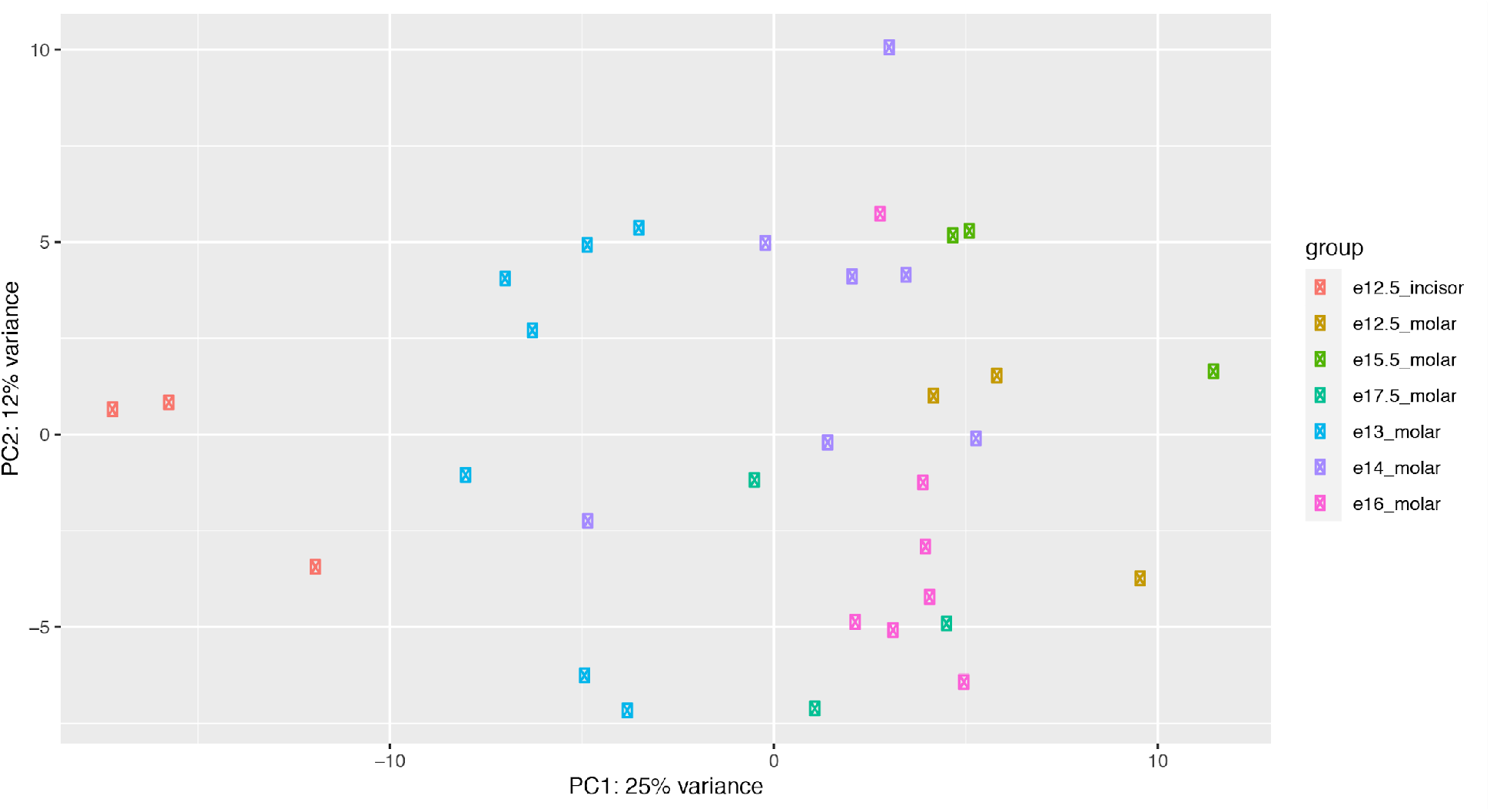
Principal Component Analysis (PCA) plot of bulk RNA-seq data obtained from [82,83], grouped by timepoint and tooth morphology. PCs determined using vst-transformed counts from RUVseq-corrected DESeq2 object.

**Supplemental Figure 5.**
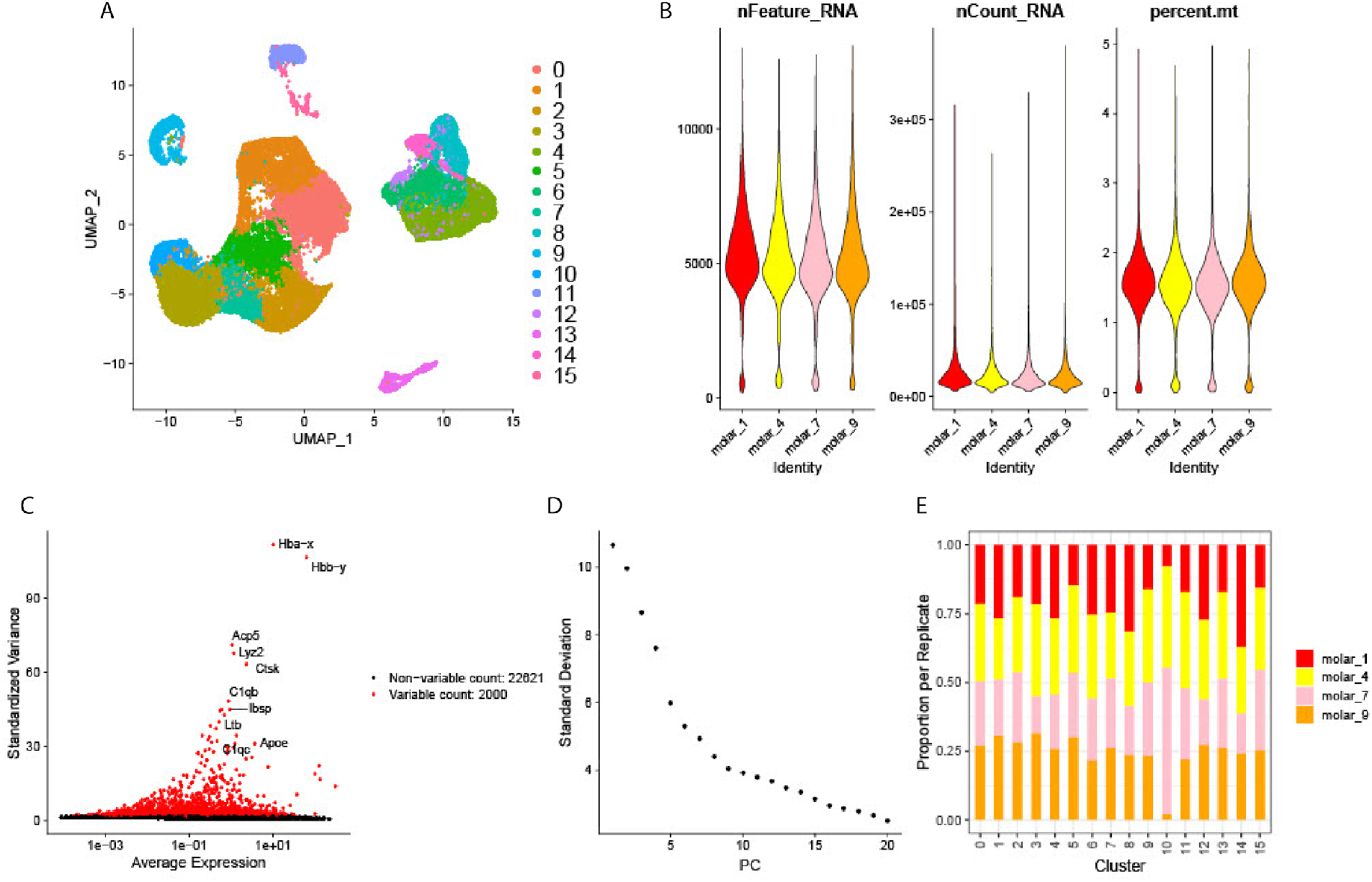
A. Initial clustered UMAP of cell states determined through Seurat analysis of 4 replicates of E14 mouse mandibular molars (n=33,886 cells) from [82]. B. Quality control of cells from A, as demonstrated by number of unique features per cell (nFeature_RNA), total number of RNA counts per cell (nCount_RNA), and percent mitochondria per cell (percent.mt), as demonstrated per replicate. C. Most highly variably expressed genes across all cells. D. Elbow plot ranking principal components of the dataset based on the percentage of variance within each PC. E. Per-replicate contribution of cells to each cluster demonstrates that cluster 10 (bone progenitor cells) is variable across replicates.

**Supplemental Figure 6.**
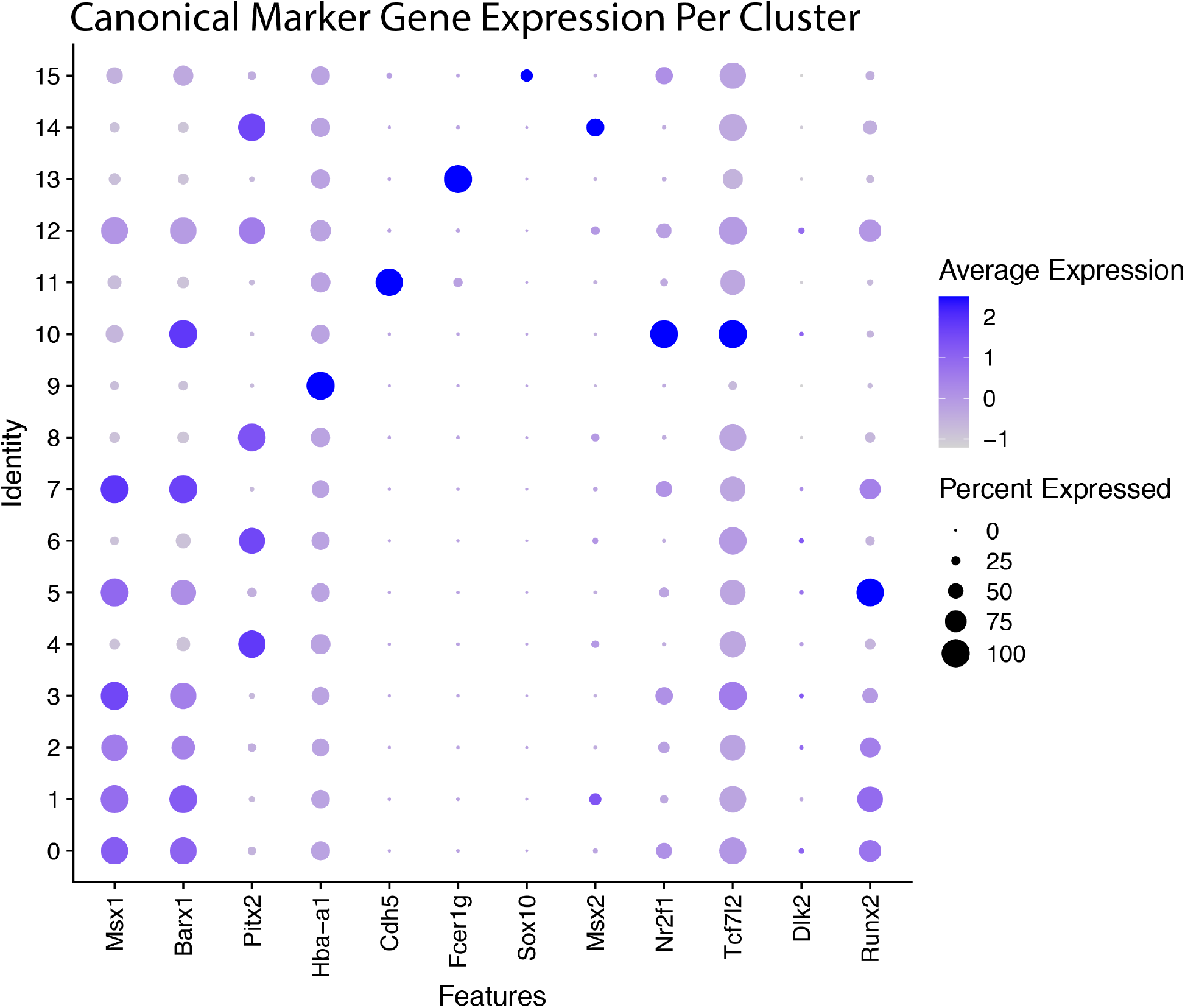
Dot plot of canonical tooth marker gene expression across all clusters from Supplemental Figure 5A. Color indicates average expression across cells per cluster, size indicates the percentage of cells per cluster which express that gene.

**Supplemental Figure 7.**
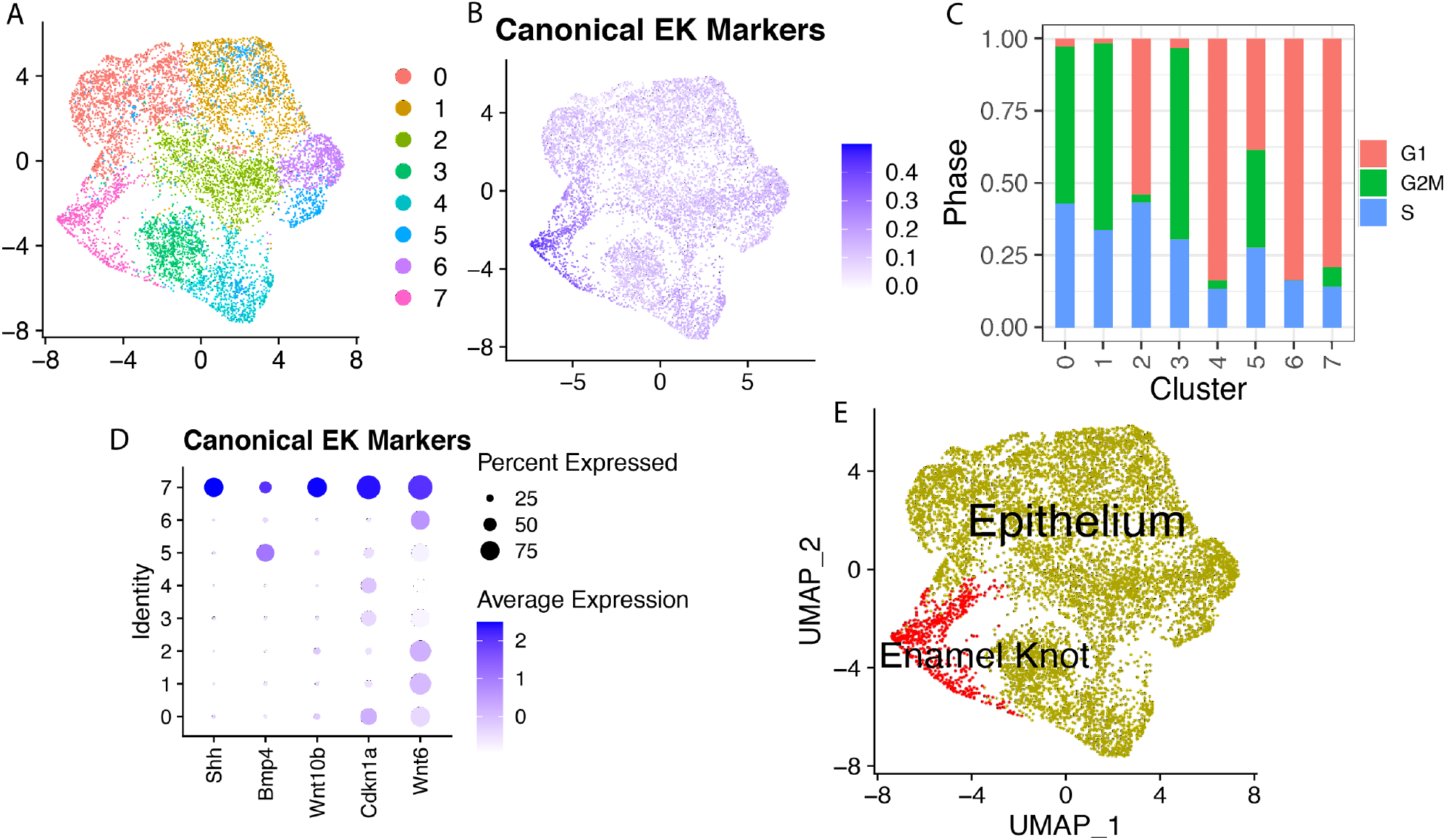
A. Initial sub clustered UMAP of cell states determined through Seurat analysis of epithelial cells from 4 replicates of E14 mouse mandibular molars (n=33,886 cells) from [82]. B. UMAP overlay of module scores of combined expression of canonical enamel knot markers, calculated by Seurat AddModuleScore. C. Predicted cell phase across epithelial cell clusters, assigned using Seurat CellCycleScoring modality. D. Dot plot of canonical EK markers across epithelial subclusters from A. Color indicates average expression across all cells per cluster, size indicates percentage of cells per cluster expressing that gene. E. Annotated UMAP from A.

**Supplemental Figure 8.**
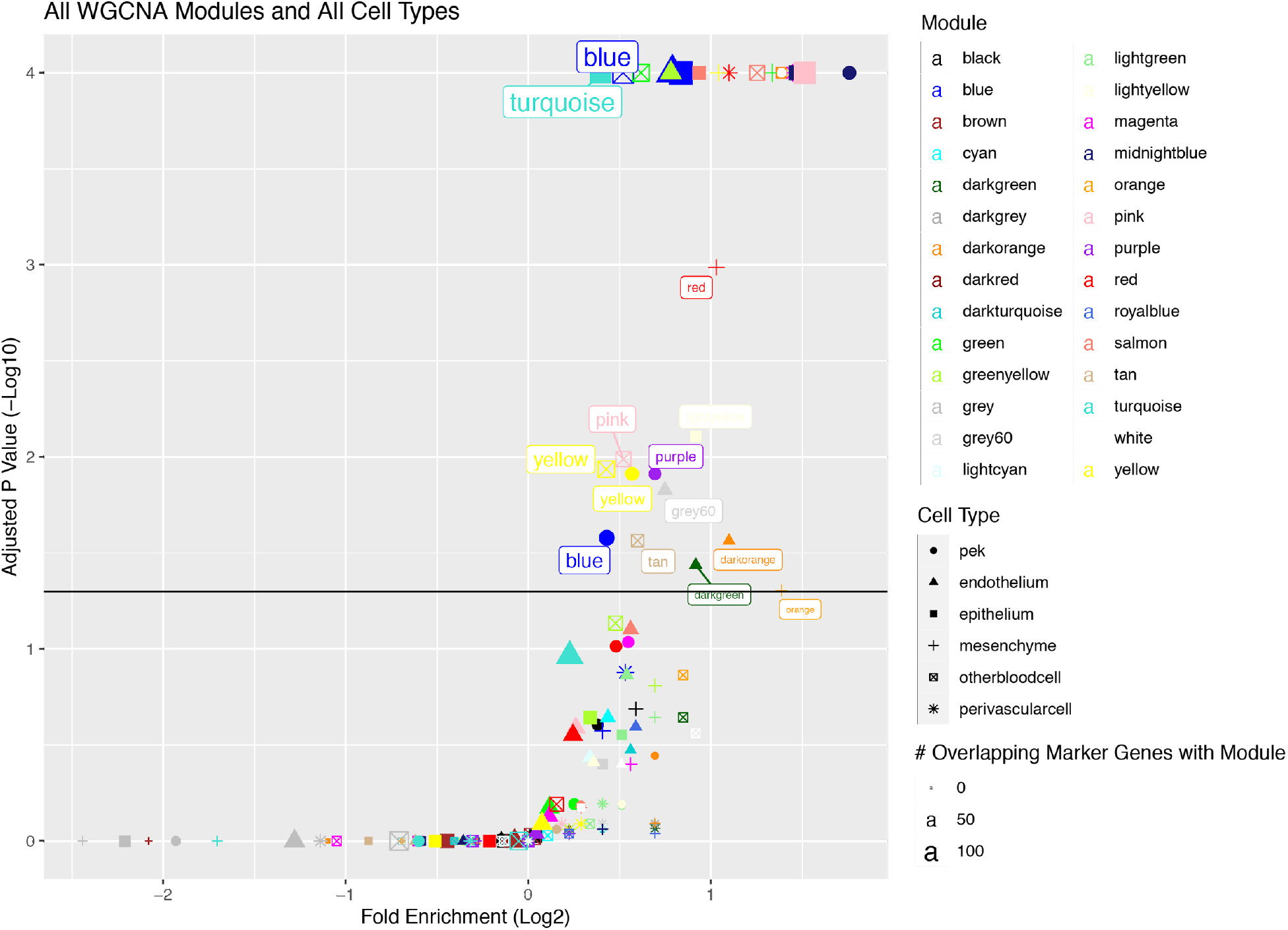
Combined enrichment findings from Figure 4E-I. Permutation enrichment analysis of marker genes from each annotated scRNA-seq cluster within genes from each WGCNA module, excepting Red Blood Cells cluster (did not show enrichment of any modules). Points are named according to the WGCNA module, and point size indicates the number of genes per module which appear as marker genes in that tissue. Shape indicates the annotated cell type and size of each point indicates the number of overlapping genes between each cell type’s transcriptomic signature and each WGCNA module. Each major cell type demonstrates enrichment of marker genes within at least 1 WGCNA module, indicating these modules are active in specific cell types of the developing tooth. The magnitude of these enrichments and their significance are comparable across cell types, demonstrated by the variety of cell types (shapes) demonstrating significant enrichment in multiple different WGCNA modules (top right points).

**Supplemental Figure 9.**
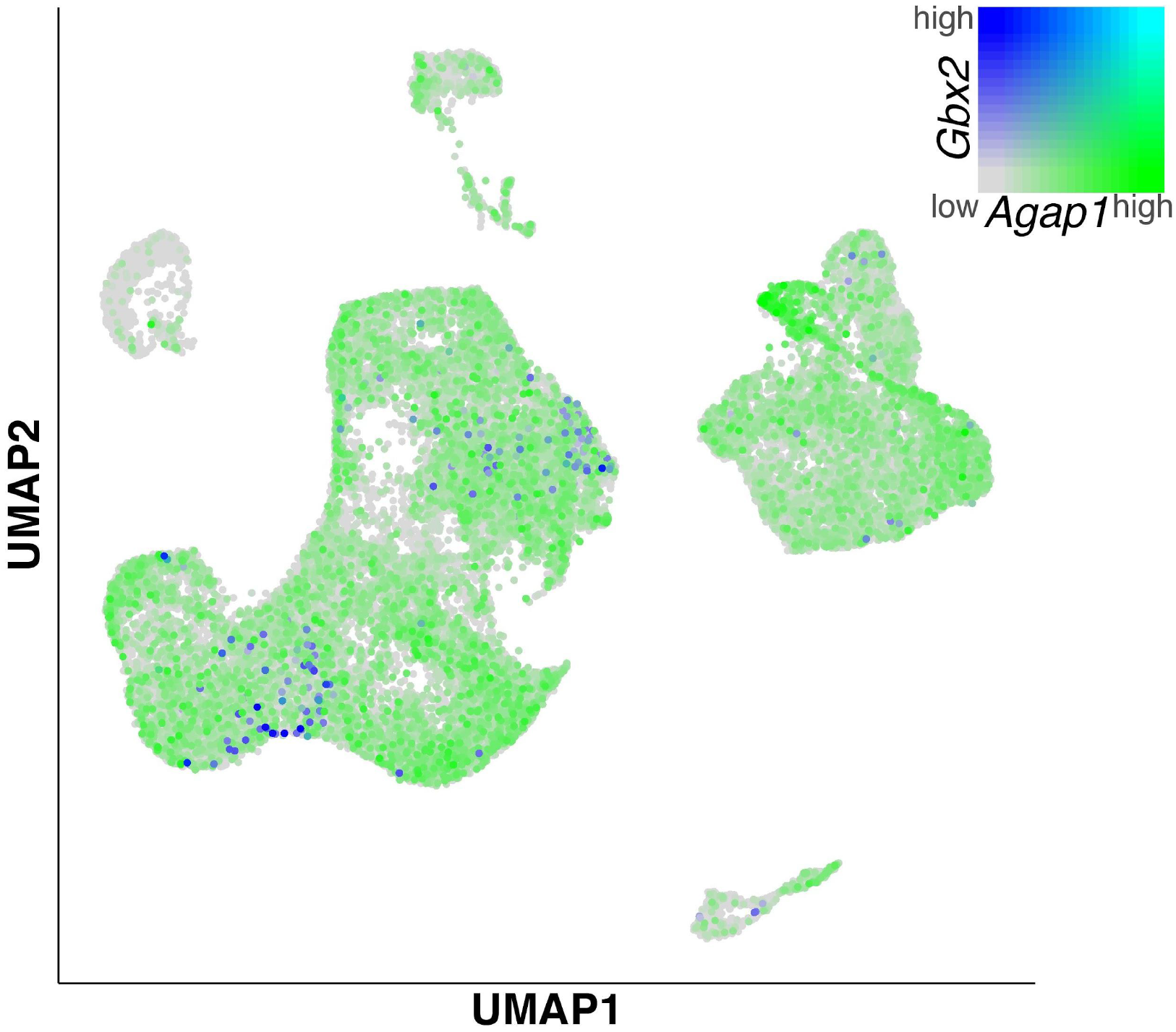
Co-Expression overlay of UMAP from A demonstrates no co-expression of *Gbx2/Agap1+* (cerulean), and the relatively low expression of *Gbx2* compared to *Agap1* in this tissue.

**Supplemental Figure 10.**
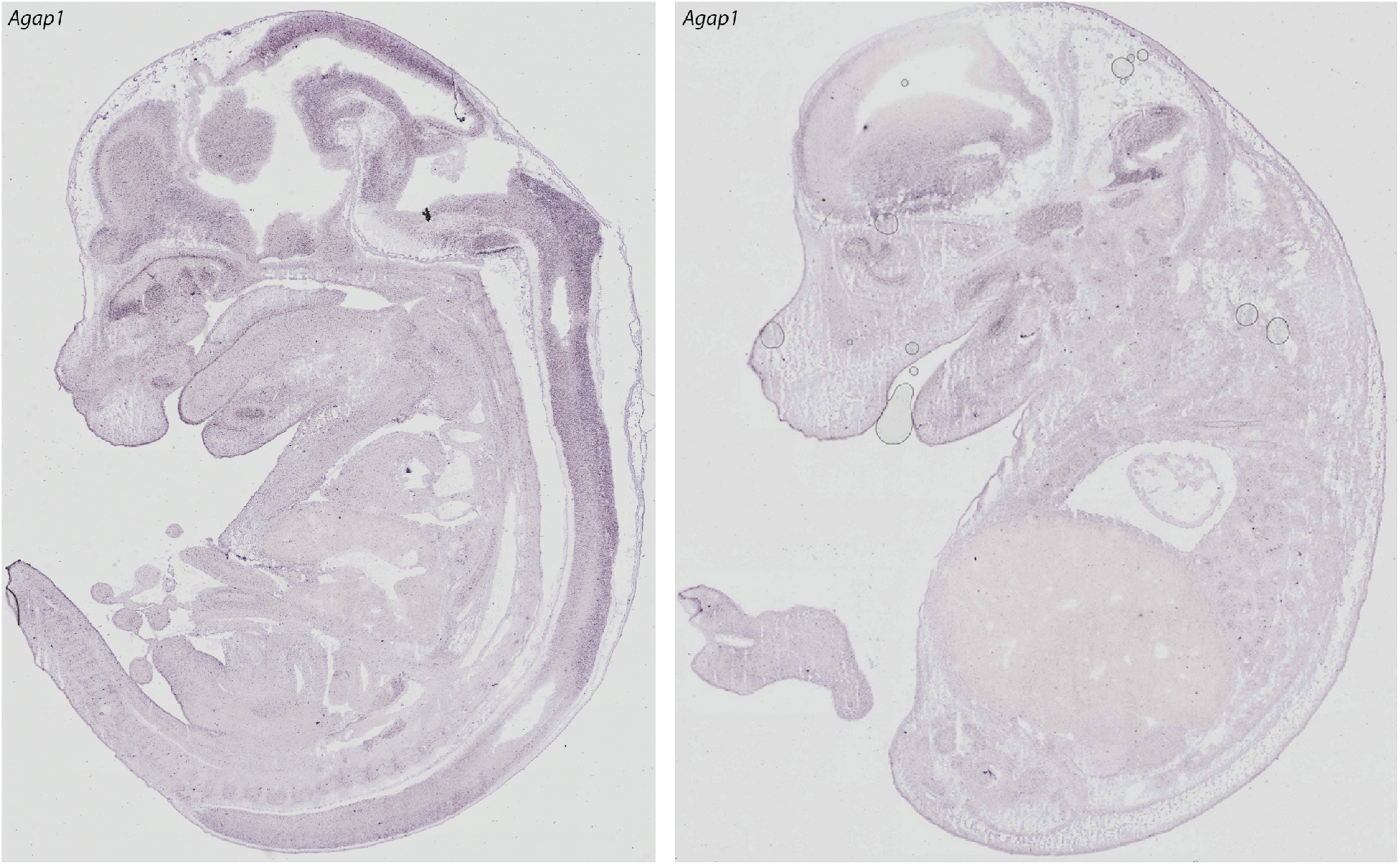
*In Situ* hybridization images of *Agap1* in E14.5 mice from GenePaint [65,87–89]. Images are expanded whole-body images from Figure 5.

**Supplemental Figure 11.**
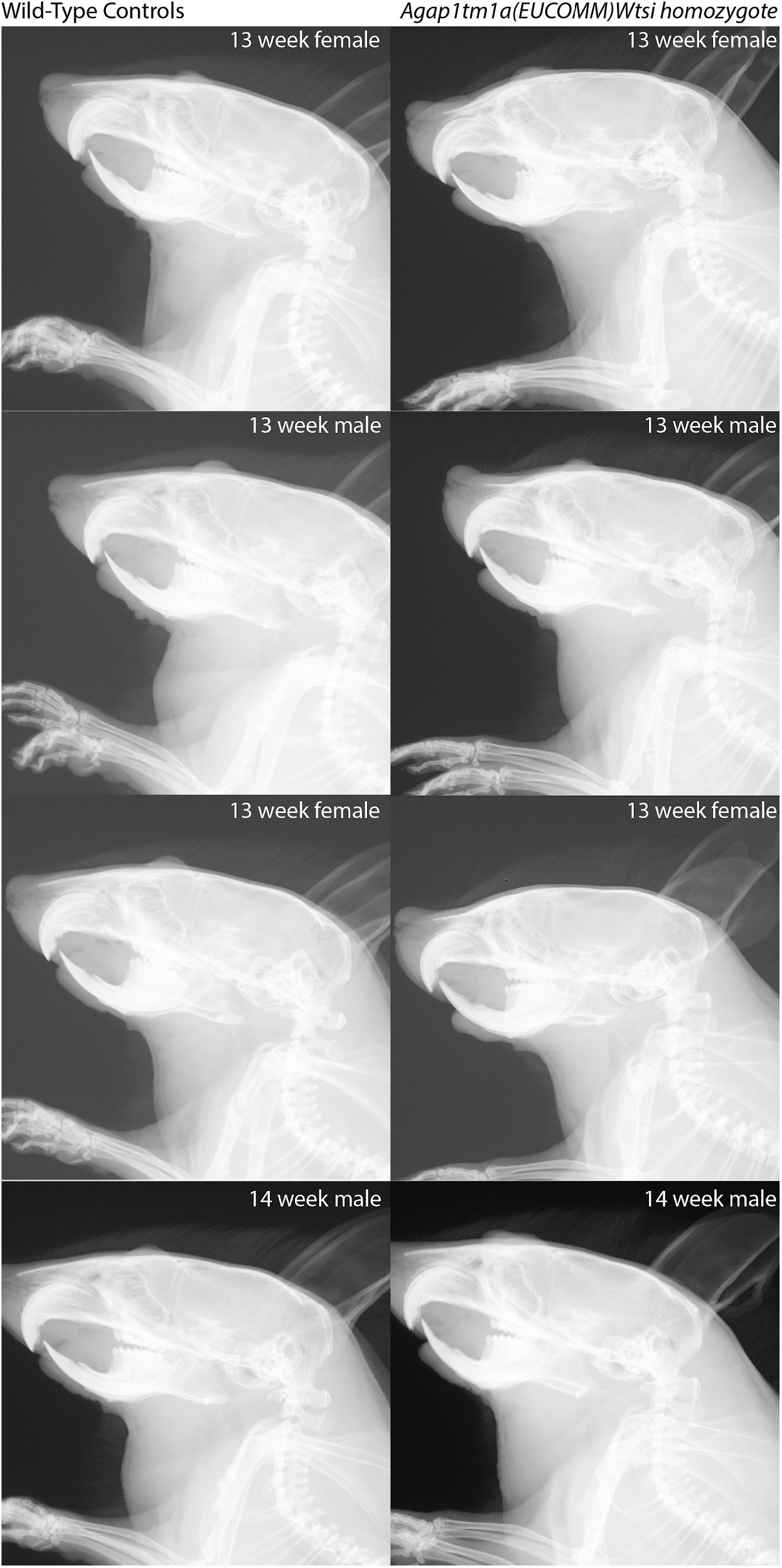
X-ray images of 13-14 week old male and female *Agap1-/-* mice and wild type, age and sex-matched littermates. Images from KOMP [96].

